# Clonal Expansion and Diversification of Germinal Center and Memory B Cell Responses to Booster Immunization in Primates

**DOI:** 10.1101/2025.06.27.661994

**Authors:** Lachlan P. Deimel, Yoshiaki Nishimura, Gabriela S. Silva Santos, Viren A. Baharani, Brianna Hernandez, Thiago Y. Oliveira, Andrew J. MacLean, Marie Canis, Sadman Shawraz, Anna Gazumyan, Harald Hartweger, Paul D. Bieniasz, Theodora Hatziioannou, Malcolm A. Martin, Michel C. Nussenzweig

## Abstract

Effective vaccines elicit B cell clonal expansion in germinal centers (GCs) that produce memory B cells and antibody secreting plasma cells. Studies in mice indicate that, whereas the plasma cell compartment is enriched for cells producing high affinity antibodies, the memory pool is more diverse and contains only a relatively small proportion of higher affinity cells. Upon boosting, murine memory B cells producing high affinity antibodies tend to develop into plasma cells but few if any re-enter GCs. However, mice live for only a few weeks in nature, and in keeping with the rather limited requirement for immune memory, this compartment comprises only 1–2% of all B cells. In contrast, memory accounts for nearly 50% of all B cells in primates. Here we examine memory and GC B cell responses in rhesus macaques immunized and boosted ipsilaterally or contralaterally with an mRNA vaccine encoding severe acute respiratory syndrome coronavirus 2 (SARS-CoV-2) Spike protein. The neutralizing activity of antibodies cloned from the memory compartment, and the size of the compartment, was independent of the site of boosting. Moreover, in primates, memory B cells enter and undergo iterative expansion in newly developing GCs when boosting is at a site distal to the site of priming. Thus, in primates, high affinity memory B cells constitute a reservoir that actively participates in further development of immunity irrespective of the anatomical site of vaccine boosting.

**Highlights:**

- Clonal overlap between primate memory and germinal center B cell compartments following booster immunization.
- Neutralization activity of the memory and germinal center compartments are independent of the boost site.
- Relationship between site of booster immunization (ipsilateral versus contralateral) and development of memory and germinal center (GC) responses in primates

## Introduction

Memory B cells are essential immune components contributing to long-term protection following vaccination and infection.^1–5^ The memory compartment develops longitudinally throughout an immune reaction, arising as export products from the GC or from activated extrafollicular B cells.^1–5^ Memory B cells exhibit strong clonal diversity and produce antibodies with a broad range of affinities, qualities that appear to help provide protection against evolving pathogens.^6–9^ Fate-mapping studies in mice demonstrate that upon re-exposure to antigen memory B cells expressing high affinity antibodies preferentially differentiate into plasma cells and rarely re-enter GCs.^6,10,11^ Instead, secondary murine GCs are predominantly seeded by naïve B cells expressing low-affinity antibodies. This phenomenon has been attributed to epitope masking by pre-existing antibodies and to changes in the memory T cell compartment.^6,10,11^

In contrast, short prime-boost intervals at the original site of immunization, or escalating dose immunization, can ‘re-fuel’ murine GC reactions by replenishing waning antigen availability.^11–14^ In mice, boosting at a distal site, or after cessation of the initial reaction, produces *de novo* GCs that recruit B cells primarily from the naïve compartment.^6,10,11^ These observations have implications for both the timing and anatomical site of vaccine booster immunizations. However, primates have a much larger and phenotypically distinct memory B cell compartment compared to the mouse and whether paradigms established in rodents can be fully extended to primates is unclear. To date, human studies show contradictory serological findings on the effects of boost site, with examples of both contralateral and ipsilateral advantages observed.^15–17^ These discrepancies remain unaccounted for but could be due to timing of vaccine boosting and assay performance. In summary the effects of booster immunization on GC and memory B cell clonal dynamics in primates remains to be defined precisely.

Here, we report on the B cell responses in a cohort of rhesus macaques who received booster vaccines at ipsilaterally or distally at a contralateral site. Complete lymph node excision and spleen biopsy recovered large numbers of GC B cells enabling analysis of cross-compartment clonal overlap and GC B cell evolution post-boost, which is not possible with fine needle biopsy. Our data indicate that clones of antigen binding memory cells in the systemic circulation may be recalled to GCs that are formed *de novo* upon contralateral boosting. Thus, the memory B cell compartment in primates comprises a pool of cells that participates in recall GC responses irrespective of the site of vaccination.

## Results

### B cell responses to booster vaccination in Rhesus macaques

To examine the development of memory B cell responses in primates and the participation of these cells in booster responses, we immunized six rhesus macaques (Table S1) with three doses of an mRNA vaccine given at intervals corresponding to the original SARS vaccination program in humans. The third dose was given either ipsilaterally or contralaterally (Figure 1A). Antibody titres in plasma were monitored longitudinally. Reactivity against Wu-1 Spike waned 4.5-fold in the 8-weeks following vax2 (Figure 1B). The highest titres were detected following the third dose, which was 2.7-fold higher than that after the second immunization (Figure 1B). As observed in murine and human studies, no differences in titres were detected between macaques boosted ipsilaterally or contralaterally (Figure 1B).^16,18,19^ We tested plasma antibody binding activity to the receptor binding domain (RBD) derived from autologous strains (Wu-1 and BA.5; constituents of the vaccine) and an archetypical escape variant (XBB1.5). At timepoints sampled, plasma from all animals displayed reactivity against the strains screened, and there were no differences in RBD reactivity associated with the site of the final boost (Figures 1C and S1).

**Figure 1:**
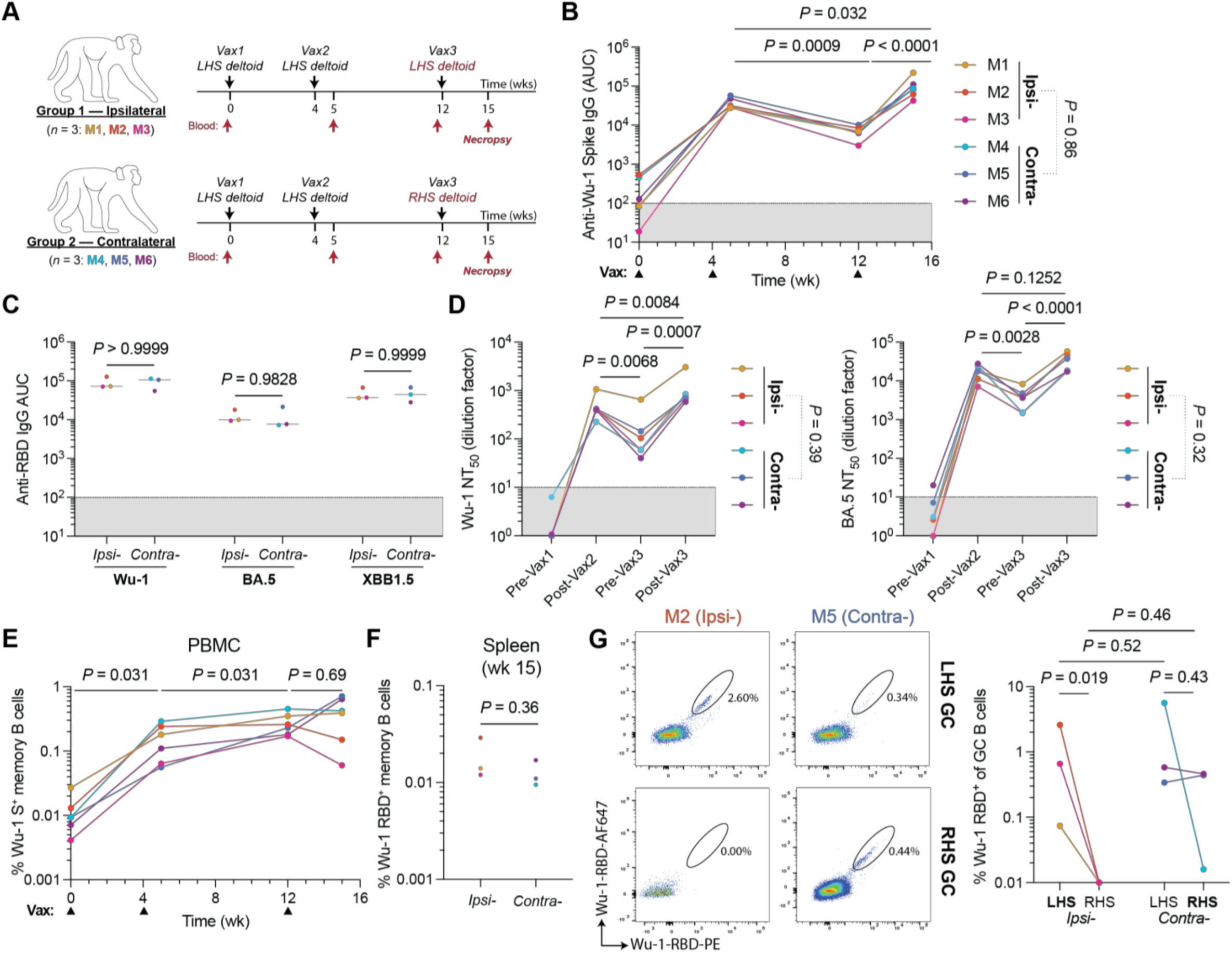
Immune responses to mRNA booster immunizations in rhesus macaques. **(A)** Schema of immunization schedules. Rhesus macaques received the Pfizer Comirnaty 2023–24 (30 µg mRNA; formulation includes Wu-1 and BA.5 Spike-encoding transcripts) vaccine. **(B)** Plasma IgG antibody titers against Wu-1 Spike over time. **(C)** Post-boost (wk 15) plasma IgG reactivity against RBD variants. Bars denote the median. **(D)** Longitudinal plasma neutralization titers (NT_50_) using pseudotyped viral particles expressing autologous Wu-1 or BA.5 whole Spike. **(E)** Percentage of whole Wu-1 Spike bait-binding circulating memory B cells (CD38^+^CD27^+^) in PBMCs. **(F)** Frequency of RBD^+^ memory B cells in the spleen. **(B, D** and **E)** Temporal pairwise comparisons were conducted using paired Tukey’s post-hoc comparisons. **(G)** Wu-1 RBD bait-binding on post-vax3 GC B cells (CD71^hi^CD38^-^) in the superior auxiliary lymph nodes at wk 15. **(B–D, E** and **G)** Terminal endpoint titers were compared between boost conditions using a Student’s t-test. Data obtained from each animal are colorized respectively, and lines conjoin datapoints from the same animal.

Plasma neutralizing activity was tested using SARS-CoV-2 Spike-pseudotyped viruses expressing Nano-luciferase.^20^ The neutralization titres against Wu-1 and BA.5 waned significantly over time between the second and third immunization (Figure 1D). Following the third immunization, these titres recovered with NT_50_ values against Wu- 1 being ∼2-fold higher compared with post-vax2 (Figure 1D).

The pre-boost circulating memory B cell compartment (CD20^+^CD38^+^CD27^+^ B cells) was enumerated by flow cytometry and their antibody sequences were documented by single cell mRNA sequencing using the 10X Chromium platform. Memory comprised an average of 43% of all circulating B cells (Figures S2A and S2B). We obtained a total of 27,787 paired heavy and light chain Ig sequences from memory B cells. Most memory cells expressed the IgG isotype, accounting for an average of 73.3% of this population, while IgM and IgA were expressed by 21.0% and 5.7% of all memory cells respectively (Figure S2C). As expected, 32–53% of the memory compartment consisted of small, expanded clones that ranged in size from 2–30 cells (Figure S2D and Table S2).

The frequency of circulating Wu-1 SARS-CoV-2 Spike binding memory B cells (CD20^+^CD38^+^CD27^+^) was quantified longitudinally (Figures 1E and S2A). Consistent with vaccination and infection in humans, Spike-binding B cells increase in the months after the second immunization in macaques to reach a maximum of 0.17–0.45% of all memory cells (Figure 1E).^21^ However, Spike specific memory B cells did not increase significantly after the third immunization (Figure 1E). Finally, the frequency of RBD- binding memory B cells were enumerated in spleen after the final boost; their frequency was similar irrespective of the site of boosting (Figures 1F, S2A and S3).

To examine the post-boost GC response in macaques, we identified GC B cells (CD38^-^ CD71^hi^; Figures S4A and S4B)^13^ in superior and inferior axillary lymph node clusters (Figure S4C). In the absence of immunization, GCs can be detected in unimmunized macaque lymph nodes, likely due to occasional breaks in skin barrier, but SARS-CoV- 2 binding B cells are absent (Figure S4D). GCs ranged from 3.5–8.3% of all B cells in the lymph nodes of the ipsilaterally boosted animals, and an average of 2-fold smaller in the unimmunized RHS lymph nodes (Figure S4E). In contrast, GC size was not significantly different between the two sides in contralaterally immunized animals, which was likely due to ongoing GC responses induced by vax1 and vax2 on the lefthand side (LHS; Figure S4E). As expected, RBD bait-binding GC B cells were found primarily in the superior auxiliary nodes that drain the sites of immunization (Figure S4F). The proportion of RBD-binding GC B cells post-boost was low, typically representing <0.5%, which is in keeping with the fraction of such cells in the memory compartment (Figures 1E, 1G, S4F and S4G). There were few if any detectable RBD binding GC B cells in lymph nodes draining regions that had not been subjected to immunization, including lymph nodes on the unimmunised side in the ipsilaterally boosted animals (Figures 1G and S4G). In 2 of the 3 contralaterally boosted animals (M5 and M6), RBD binding cells were present in similar numbers in both arms (Figures 1G and S4G). The one exception, M4 had the smallest GCs and showed a lower fraction of RBD bait-binding GC B cells in the contralaterally boosted lymph nodes than what remained in the site of the original immunization (Figures 1G and S4G).

### GC responses after multiple immunizations

The relatively small GC size and low frequency of RBD binding cells in macaques were somewhat unexpected. To determine whether these observations are primate specific, we performed parallel experiments in C57BL/6 mice. Mice were immunized with 0.5 µg Wu-1 SARS-CoV-2 Spike mRNA, with the third dose given either ipsilaterally or contralaterally (Figure 2A). Irrespective of site of the final booster, GCs were smaller after the third immunization compared to the second (Figure 2D and S5A). Similar reduction in post-vax3 GC size was also seen in mice immunized with 2 µg recombinant ovalbumin (OVA) in alhydrogel (Figure S5B). In addition to the reduced GC size post-vax3, an average of only ∼0.5% of the B cells in the GCs bound to RBD, irrespective of the booster site (Figure 2E). We conclude that in immunized mice and macaques, the small GCs elicited by the third immunization contains only a small proportion of antigen binding cells.

**Figure 2:**
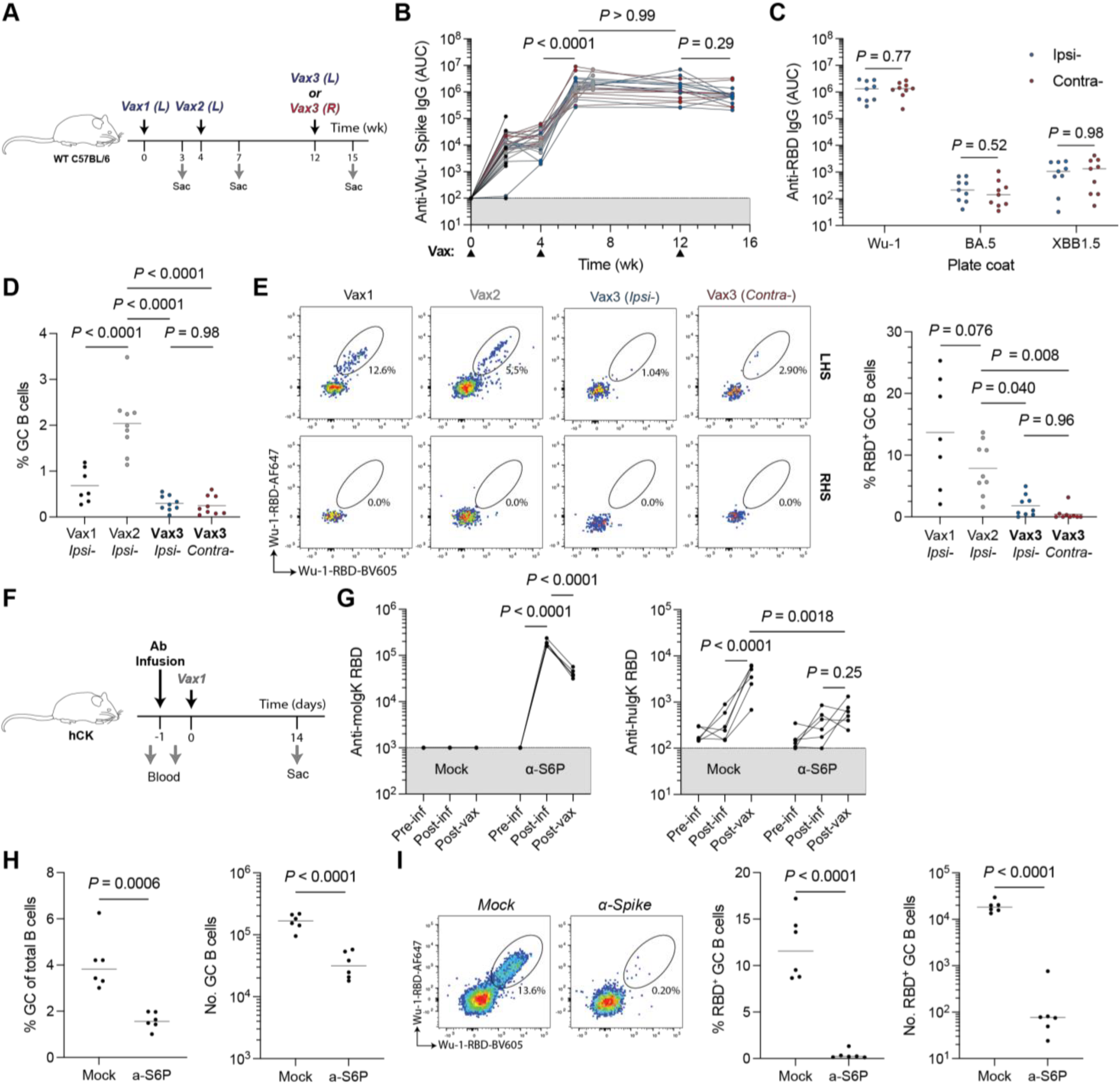
Pre-existing serology affects germinal center reactions to booster immunizations. **(A)** Experimental scheme for **(B–E)**. Mice were immunized with 0.5 µg mRNA encoding Wu-1 SARS-CoV-2 Spike protein in LNPs. Immunizations were administered into the gastrocnemius muscle. Data are pooled from two independent experiments. **(B)** Longitudinal serum antibody IgG titres against the autologous Wu-1 SARS-CoV-2 Spike. Connected dots denote titers from the same animal at different timepoints. **(C)** Serum antibody titers against RBD variants at the terminal (wk 15) timepoint. Groups compared using student’s t-test. **(E)** The percentage of GC B cells (Fas^+^CD38^-^) in lymph nodes after each immunization. **(F)** Experimental scheme for **(G–I)**. Animals received intravenous infusions of 1 mg purified serum Ig from naïve (mock) or SARS-CoV-2 Spike-immunized donor wild-type animals. **(G)** Exogenous (mouse IgK, left) and endogenous (human IgK, right) antibody titres against Wu-1 RBD. **(H)** Germinal center size 14 days post-immunization. **(I)** Wu-1 RBD antigen-binding B cells in the GC. **(B, D, E and G),** Tukey’s two-tailed post-hoc multiple comparison was used for pairwise analyses. Comparisons in **(B and G)** are paired. **(C, H and I)** Student’s T-test was used for pairwise comparisons between groups.

To examine the impact of pre-existing antibodies on dampened GC reactions post-boost we purified IgG from naïve or SARS-CoV-2 immunized C57BL/6 mice and passively transferred it to naïve mice (Figure 2F and 2G (left)). We used human IgK- bearing mice (hCK) mice as recipients to distinguish the endogenous antibody response from passively transferred antibodies. Passive antibody infusion significantly blunted the endogenous serological response and reduced GC size and the proportion of RBD bait-binding cells (Figures 2G-2I). Thus, high titers of pre-existing antibodies restrict serologic and GC responses to booster vaccines and are consistent with earlier epitope masking studies in both monoclonal and polyclonal settings.^11,22,23^ This does not preclude other factors, such as altered T cell help or antigen form/deposition, from also playing a role in humoral responses to repeat immunization.

### Shared B cell clones in memory and GC compartments

To evaluate the clonal dynamics between circulating memory and GC B cells in primates we obtained 493 and 463 paired Ig heavy and light chain sequences from macaque single RBD binding cells from lymph node GC, and splenic memory B cells respectively after the last boost (Table S3). We compared the V_H_-gene segment usage between the pooled single RBD binding B cells and bulk memory repertoire (Figure S2D and Table S2). Clones utilising *IGHV4-79*, *IGHV5-157*, *IGHV4-AFL* and *IGHV4- NL_38* gene segments were over-represented among the RBD binding memory antibodies from most animals (Figure S6A).

The Ig clonal overlap between the memory compartment and post-vax3 GC was evaluated to understand the relationship between the 2 compartments. We identified cross-compartment clonal overlap in all animals (Figure 3A and S6B). The proportion of RBD binding clones shared between the GC and memory compartments was variable, ranging from 10–50% and was independent of the boosting site (*P* = 0.18; Figure S6C). Like observations in mice,^2,3^ macaque memory cells were clonally more diverse than the GC B cell compartment irrespective of boost side (*P* < 0.0001; Figure S6D). Finally, there was no difference in clonal diversity between GCs on the two sides in the contralaterally boosted animals (Figure S6D).

**Figure 3:**
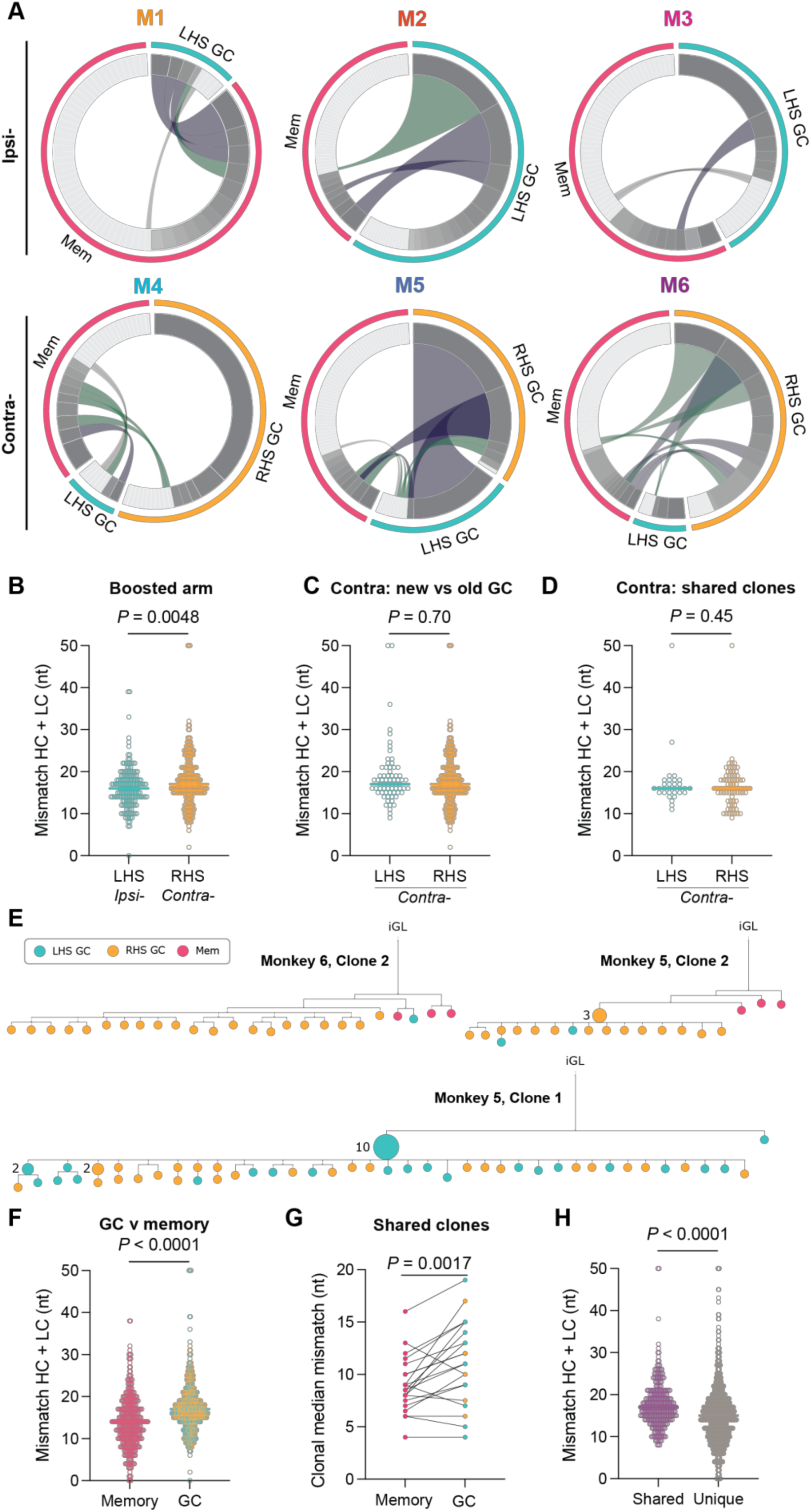
B cell clones in the germinal center and memory compartments following booster immunizations. **(A)** Circos plots showing B cell clone sharing between Wu-1 RBD bait-binding memory and GC B cells in Rhesus macaques following three mRNA SARS-CoV-2 immunizations. Segment size is proportional to the clone size, with the small light grey- colored segments denoting singlets and the darker grey indicating expanded clones. The magenta wedge denotes splenic memory clones, teal denotes GC B cell clones identified in the LHS auxiliary lymph nodes (aLN) and yellow denotes GC B cell clones identified in the RHS aLN. Conjoined cross-compartment segments indicate clones present in multiple compartments; grey = singlet–singlet, green = expanded clone–singlet, and purple = expanded clone–expanded clone. **(B)** Germline nucleotide mismatch in GC B cells from the boosted side from each site condition. **(C and D) (C)** Graph of the nucleotide mismatch in both sides in contralaterally-boosted macaques and **(D)** SHM of clones belonging to families present in both the LHS and RHS GC. **(E)** Phylogenetic trees of individual B cell clonal families based on Ig heavy and light sequences rooted to their inferred germline precursor (iGL). Nodes comprise of a single cell unless otherwise marked. Node color denotes the location and fate of the originating cell. **(F)** Plots showing the germline nucleotide mismatch of heavy chain and light chain sequences of memory and GC B cells. **(G)** Plot showing the paired germline nucleotide mismatch of clones with family members present in both the GC and memory compartment. In cases where more than one cell is in a compartment, the median was used. Data were compared using a paired Student’s t-test. **(H)** Plot showing the germline nucleotide mismatch of cells belonging to clonal families shared across multiple compartments or only present in a single compartment. **(B–D, G and H)** Data were compared using a Student’s t-test. Each point is the mutation load (cumulative germline mismatch of heavy and light chains) for a single cell. Data are pooled from all animals.

Ipsilaterally boosted animals showed no RBD binding GC B cells in the contralateral side and therefore no overlap between the 2 sides. Thus, immune responses to the mRNA immunogen did not become systemic and were anatomically restricted. GCs in contralaterally boosted animals that displayed larger GCs and a greater number of RBD binding B cells, M5 and M6, contained B cell clones that were shared between the contralaterally boosted RHS GC and the persistent LHS (Figure 3A). M5 exhibited 98% overlap between RBD binding clones on the two sides (Figure 3A). In conclusion, antigen binding memory B cells undergo strong proliferative expansion in recall GCs in primates.

RBD binding B cells in GCs on the boosted side had similar mutation loads, despite reaching statistical significance (Median: 17 (ipsi-) and 18 nt (contra-); *P* = 0.0048; Figure 3B). Moreover, there was no difference in SHM between RBD binding GC B cells in LHS and RHS lymph nodes of the contralaterally-boosted animals (Figure 3C and 3D). Although this finding may be counter-intuitive because GCs on the LHS have persisted for several months with more time to acquire mutations, this phenomenon may be accounted for by progressive GC clonal turn-over and influx of newly activated B cells.^24,25^

To examine the relationship between memory and GC B cell clones we constructed phylogenetic trees based on their combined Ig heavy and light chain nucleotide sequences (Figure 3E and S7). Individual memory B cells expressed antibodies found throughout the phylogenetic trees, in both discrete clades absent from GCs (Figure 3E, M6-C2 and M5-C2) and embedded amongst GC B cells (Figure S7, M2-C1 and M6-C3). In addition, we found large clones of identical expanded cells^26,27^ with subsequent diversification in GCs on both sides in contralaterally boosted animals (Figure 3E, M5-C1). As might be expected from earlier export, RBD binding memory B cells were generally less mutated than their GC counterparts that continued undergoing SHM (Figure 3F–3H).

We cloned and expressed 243 monoclonal antibodies from RBD binding B cells from all six animals. This set was comprised of at least one member from each clonally expanded and/or compartment shared family, as well as a random subset of singlets of both memory and GC origins (Table S4). Of the selected antibodies, 230 (95%) bound to at least one RBD variant with an EC_50_ < 5 µg/mL in ELISA, indicating efficient and specific B cell isolation (Figure S8A). Antibodies bound Wu-1 RBD with a mean EC_50_ of 27.5 ng/mL and a median of 13.8 ng/mL (Figure S8A). We examined the binding breadth of the monoclonal antibodies by screening for reactivity to BA.5 and XBB1.5, and the highly divergent SARS-CoV1 RBDs. The mean EC_50_ of BA.5 was higher than against Wu-1, while clones reacted with XBB1.5 with a lower average EC_50_ (Figure S8A). The reactivity of antibodies against SARS-CoV-1 was low, with only 63 (26%) exhibiting an EC_50_ < 5 µg/mL. Approximately 24% of binding clones were reactive with all four strains screened (Figure S8B).

To examine whether the booster site affected the circulating memory B cell binding activity and breadth, we compared memory-derived monoclonal antibody reactivity (Figure S8C). We found no significant difference in EC_50_ values or breadth of binding activity between antibodies cloned from the memory compartment or GCs of animals boosted ipsilaterally or contralaterally (Figure S8C and S8D). In the contralaterally- boosted animals, the antibodies cloned from RBD binding GC B cells from the LHS and RHS were essentially indistinguishable in activity and breadth (Figure S8D). Binding breadth between the memory and GC B cell-derived antibodies was equivalent (*P* = 0.77; Figure S8E).

A subset of 150 monoclonal antibodies that bound RBD by ELISA were tested for neutralization activity against pseudotyped viruses expressing SARS-CoV-2 Spike derived from the autologous Wu-1 and BA.5 strains, as well as prototypical escape variants XBB1.5 and JN.1 (Figure 4).^28^ Of the antibodies tested, 98 (65.5%), neutralized Wu-1, with a mean NT_50_ of 1.58 µg/mL (Figure 4A). Consistent with the values obtained from plasma, neutralizing activity was better against BA.5 than Wu-1 (Figure 1D and Figure 4A, *P* = 0.0074). Neutralizing activity against XBB1.5 was similar to BA.5, while neutralizing activity against JN.1 resembled Wu-1 (Figure 4A).

**Figure 4:**
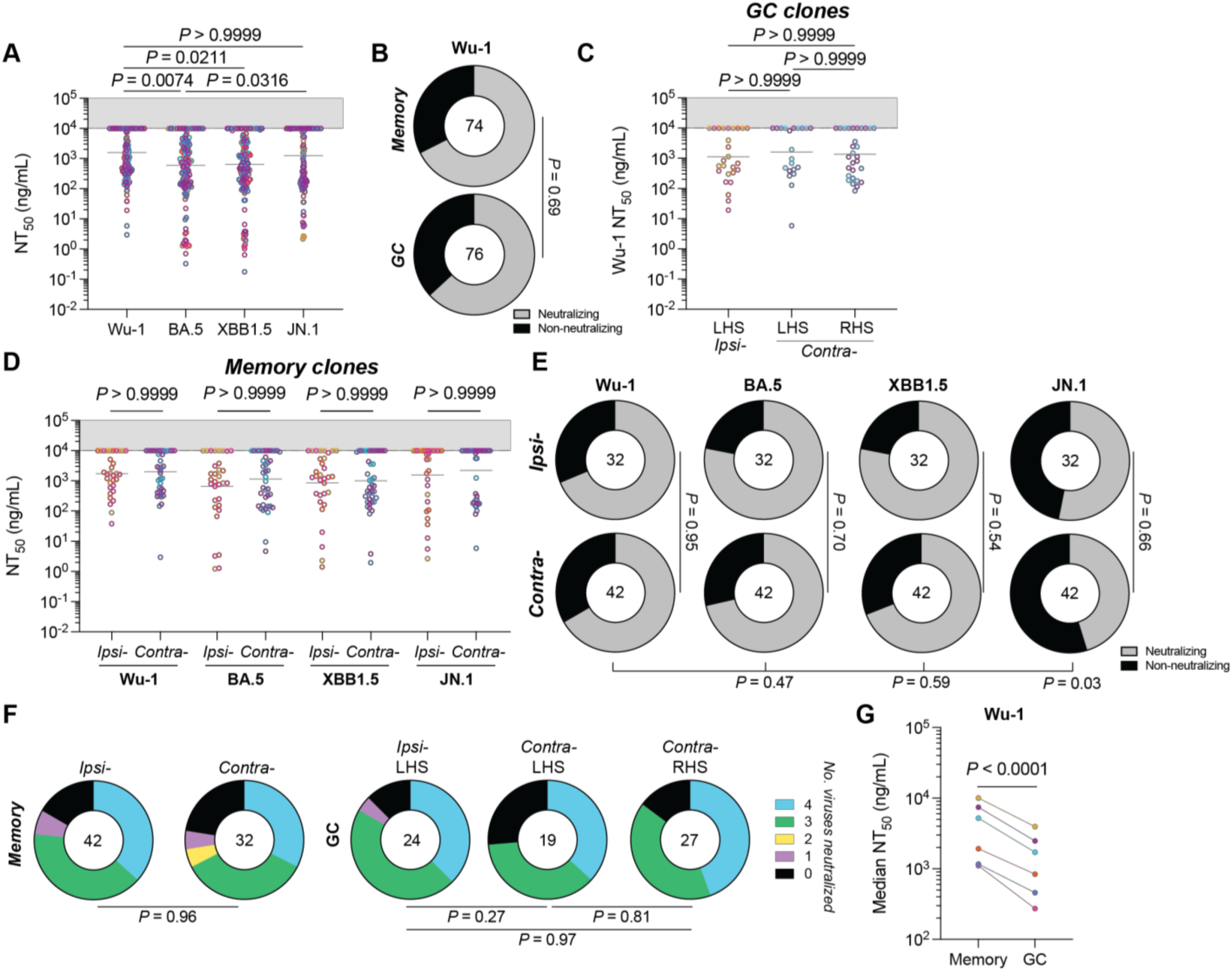
Neutralization by anti-SARS-CoV-2 RBD monoclonal antibodies. **(A)** Plot of the neutralization titers of antibodies cloned from macaque Wu-1 RBD bait-binding memory and GC B cells against autologous (Wu-1 and BA.5) and prototypical escape variants (XBB1.5 and JN.1). **(B)** Donut plots showing the proportion of antibodies deemed non-neutralizing (NT_50_ > 10 µg/mL) against Wu-1. **(C)** Graph showing Wu-1 neutralization titers of antibodies cloned from GC B cells. Antibodies were partitioned with respect to their originating clone boost site and node side. **(A and C)** Data were compared using a Kruskal-Wallis test. **(D and E) (D)** Graph showing the NT_50_ values of antibodies cloned from memory B cells from either ipsilaterally or contralaterally boosted animals and **(E)** donut plots of memory-derived antibodies categorized as neutralizing or non- neutralizing. **(F)** Donut plots showing the number of viruses neutralized by antibodies derived from either memory or GC B cell compartments. **(A, C** and **D)** Dots show the NT_50_ value from a single monoclonal antibody and are colorized according to animal the clone was isolated from. Bars denote the geometric mean. **(B, E** and **F)** Data were compared using chi-square tests, applying a Benjamini-Hochberg *P*-value correction where multiple pairwise comparisons were conducted. **(G)** Plot showing the paired average Wu-1 NT_50_ among antibodies cloned from memory or GC B cells. Data were compared using a paired Student’s t-test.

When compared directly, the fraction of neutralizing and non-neutralizing monoclonals obtained from RBD binding B cells from GC and memory was similar (P = 0.69; Figure 4B). Moreover, the site of boosting did not affect the neutralizing activity or breadth of the antibodies obtained from either the GC or memory (Figure 4C–4F). The data suggest that at a clonal population level, the booster site does not detectably affect the memory pool or the composition of antigen binding B cells in GCs or the neutralizing activity of the antibodies they produce.

The overall neutralization potency of antibodies obtained from GCs was ∼2.9-fold higher than memory (*P* < 0.0001; Figure 4G). However, the neutralization titres of memory and GC B cells of the same clonal family were highly variable, with instances of GC clones improving, staying the same or worsening compared to their corresponding memory clones (Figure 5A–4D and S9). Finally, shared GC and memory clones showed no difference in neutralization breadth or potency when compared to clones found only in GCs (Figure 5E).

**Figure 5:**
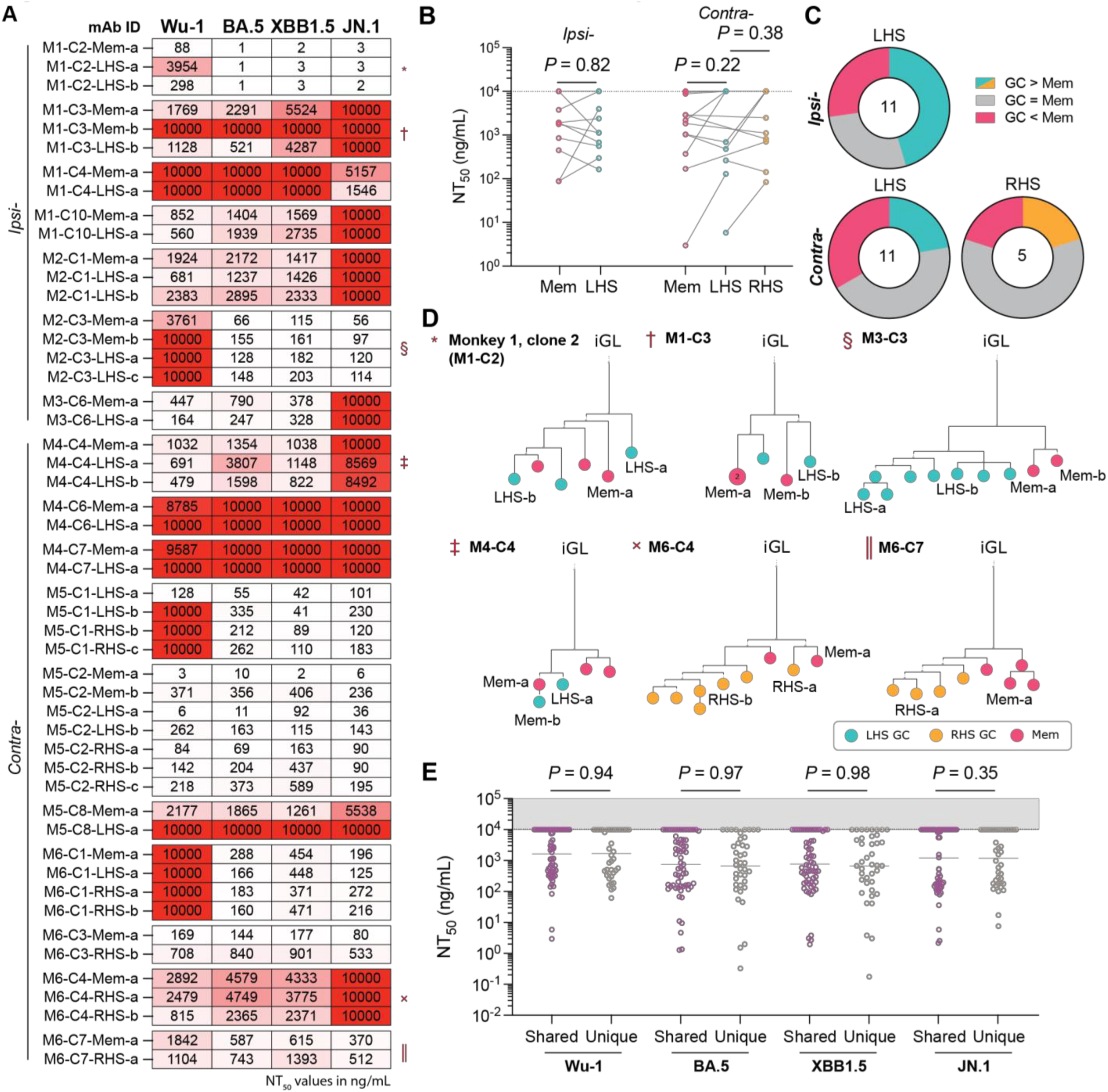
Clonal variation in neutralization between memory and GC B cells. **(A)** Heatmap showing the NT_50_ values of monoclonal antibodies derived from B cells whose clonal family was represented in multiple compartments. mAb ID assignments correspond to the monkey (M*x*), clone number (C*y*), origin (memory, LHS/RHS GC) and cell identifier (a,b, etc.). **(B)** Plot showing the paired Wu-1 NT_50_ values of clones shared between multiple compartments. Data compared via paired Student’s t-tests. **(C)** Donut plots showing the proportion of GC–memory pairs that neutralize Wu-1 better, worse, or approximately the same (NT_50_ within ± 25% of the highest value) as each other. **(D)** Example trees of clonal phyla where neutralization potency was tested across multiple nodes. Trees are rooted to the inferred germline revertant (iGL), and node colors connote the fate and site of the originating B cell. Nodes represent a single cell, unless otherwise marked. Symbols aside the monkey and clone IDs correspond with the respective clone segment in **(A)**. Nodes where the neutralization was tested are labelled. **(E)** Plot showing the neutralization titres of GC B cells partitioned according to whether a corresponding memory B cell was identified (shared) or whether the clone was only identified in the GC (unique). Dots represent the NT_50_ value of a single monoclonal antibody and bars denote the geometric mean. Data were compared using Mann-Whitney tests.

## Discussion

Experiments in mice indicate that high affinity antibody producing memory B cells are rarely recalled to GCs^6,10,11^ and predominantly develop into plasmablasts upon secondary challenge.^29–32^ However, mice have a much smaller memory compartment than primates.^2,3^ Thus, murine models of limited memory-to-GC re-entry may be discordant with macaques and humans, where the memory compartment is ∼25–50- fold larger and circulates systemically.^1,2,33^ Here, we describe the clonal dynamics between post-boost memory and GC compartments in a cohort of rhesus macaques.

Consistent with the larger compartment, primate memory B cells are recruited to and undergo clonal expansion in newly elicited GCs. This is typified by expanded clones of B cells found on both sides of the contralaterally boosted animals. Human studies indicate that long lived GCs produced after mRNA vaccination continuously contribute to memory and plasma cell compartments.^34^ Therefore, GC recycling of the circulating memory B cell repertoire following successive immunization is likely to be a key mechanism for expanding or maintaining existing high-affinity clones. As in mouse studies, highly mutated memory B cells differentiate into plasma cells upon boosting in humans.^35^ Li et al. studied individuals that received 2 doses of the mRNA vaccine and found that newly recruited naïve cells undergo affinity maturation following the booster, whereas highly mutated cells in circulation do not.^35^ However, individuals followed over a 12 month period that included 3 vaccine doses showed increasing levels of somatic mutation over time that was correlated with increased binding and neutralizing activity. ^21^ In contrast to prior work examinig B cells in circulation or by fine needle biopsy, our LN excisional and spleen biopsy data directly demonstrates that primate memory B cells recycle into the GC.

The circulating antibodies and the memory B cell pool produced by booster immunization are equivalent in terms of antigen binding and neutralization irrespective of booster site—that is, whether ‘re-fuelling’ persistent GCs or creating new ones. However, continued booster vaccines in primates and mice result in increasingly smaller GCs occupied by very modest numbers of high affinity antigen binding B cells independently of the site of immunization. This effect is due in part to epitope masking. ^11,22,23^ Use of heterologous booster immunizations may reduce the effects of epitope masking and raise *de novo* clones against variant epitopes, a finding supported by recent human studies.^36^

Epitope masking after multiple boosters favour B cells expressing antibodies that target subdominant non-neutralizing epitopes, or express low affinity antibodies. ^11,22,23,37,38^ Our data indicate that GCs elicited by repeat immunization expand and further diversify the B cell response, but may not further improve binding affinity or neutralization potency on the level of individual clones. This effect may be responsible for the reported increase in susceptibility to Influenza after repeated vaccination when compared to a single vaccine dose. ^39,40^ It may also explain why vaccination against HIV-1 by sequential immunization is particularly challenging.

### Limitations of the study

An important limitation of our study is the number of macaques used which impacts the sensitivity and generalizability of serological comparisons between groups. Moreover, we were unable to perform longitudinal sampling due to logistical and technical limitations. In addition, the data does not allow us to distinguish clonal re-entry in the ipsilaterally-boosted macaques. Finally, we selected B cells according to their capacity to bind RBD bait; this biases in favour of clones that are relatively high affinity and bind to a site that contains more neutralizing epitopes than other Spike domains.

## Supplemental Information

Supplementary data are provided Figures S1–S9 and Tables S1–S5.

## Acknowledgments

We thank members of the Nussenzweig, Martin, Hatziioannou and Bieniasz labs for discussions, T. Eisenreich for animal husbandry, K.-H. Yao and the clinical processing team for tissue processing, J.-P. Truman and K. Gordon for flow cytometry, J. Woodruff and H. Duan for NGS preparation, and P. Zhou, P. Laffont and K. Lenart for assistance in plasmid preparation. We also thank R. Petros, A. Bruce, E. Stanford, K. Worku, M. Young, and T. Thomas for their diligent assistance in maintaining the animals and performing procedures. We are grateful to S. Crotty for sharing their private macaque germline Ig gene segment database, and H. Sutton for coding advice.

This work was supported in part by National Institutes of Health (NIH) grant 5R37 AI037526, NIH Center for HIV/AIDS Vaccine Immunology and Immunogen Discovery (CHAVID) 1UM1AI144462-01 to M.C.N., and the Stavros Niarchos Foundation Institute for Global Infectious Disease Research. The work was also supported in part by Division of Intramural Research Program NIAD/NIH. M.C.N. is a Howard Hughes Medical Institute (HHMI) investigator.

## Declaration of interests

M.C.N. is a member of the Celldex Scientific Advisory Board.

## Author contributions

Conceptualization of project: L.P.D., M.C.N.

Methodology: L.P.D., G.S.S., V.A.B., T.Y.O., A.G., P.D.B., T.H., M.C.N.

Investigation: L.P.D., Y.N., G.S.S., V.A.B., B.H., T.Y.O., A.J.M., M.C., S.S., A.G., H.H.

Funding acquisition: P.D.B., T.H., M.A.M., M.C.N.

Project administration: P.D.B., T.H., M.A.M., M.C.N.

Supervision: H.H., P.D.B., T.H., M.A.M., M.C.N.

Writing – original draft: L.P.D., M.C.N. Writing – review & editing: L.P.D., M.C.N.

## STAR METHODS

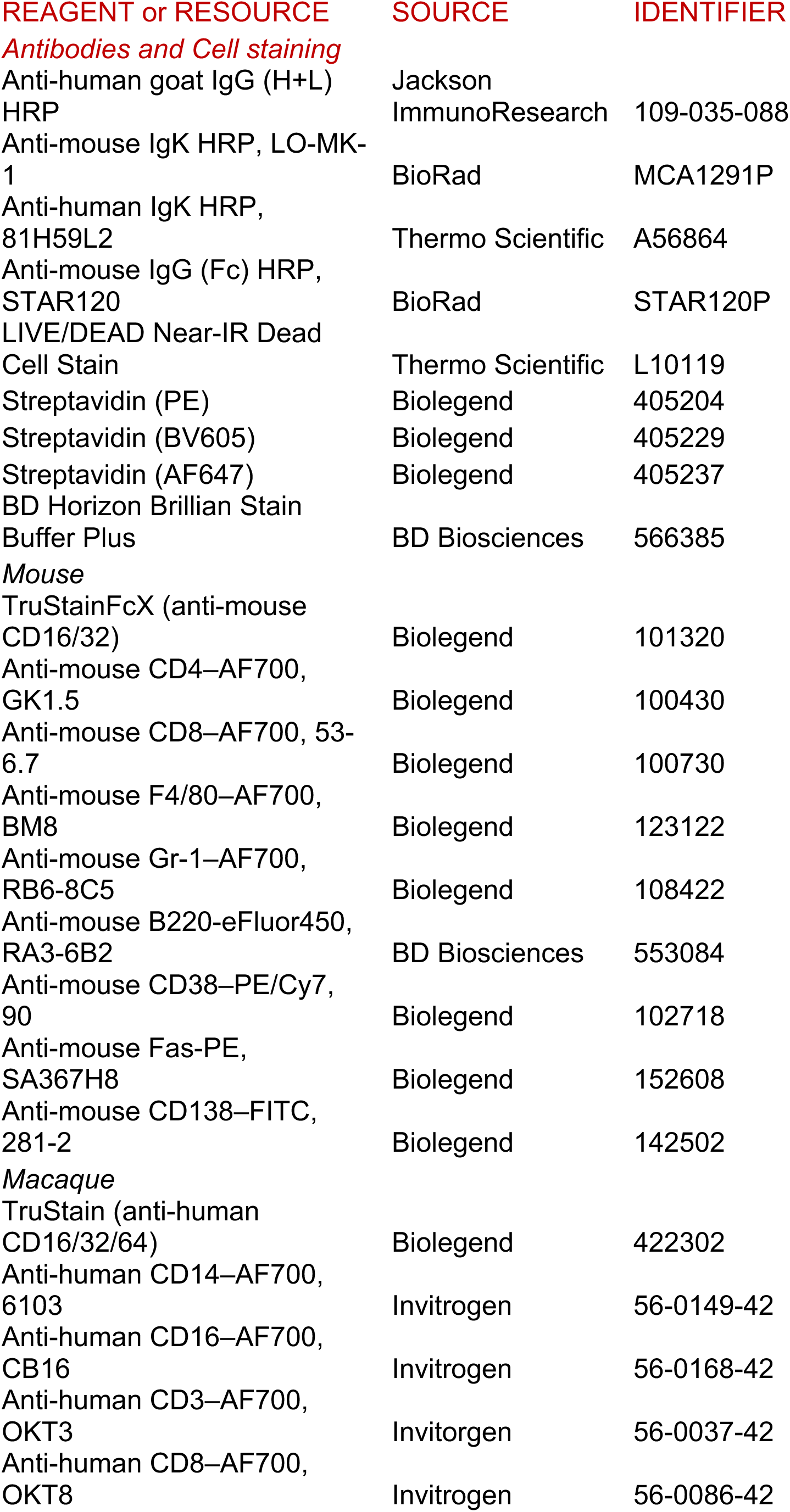
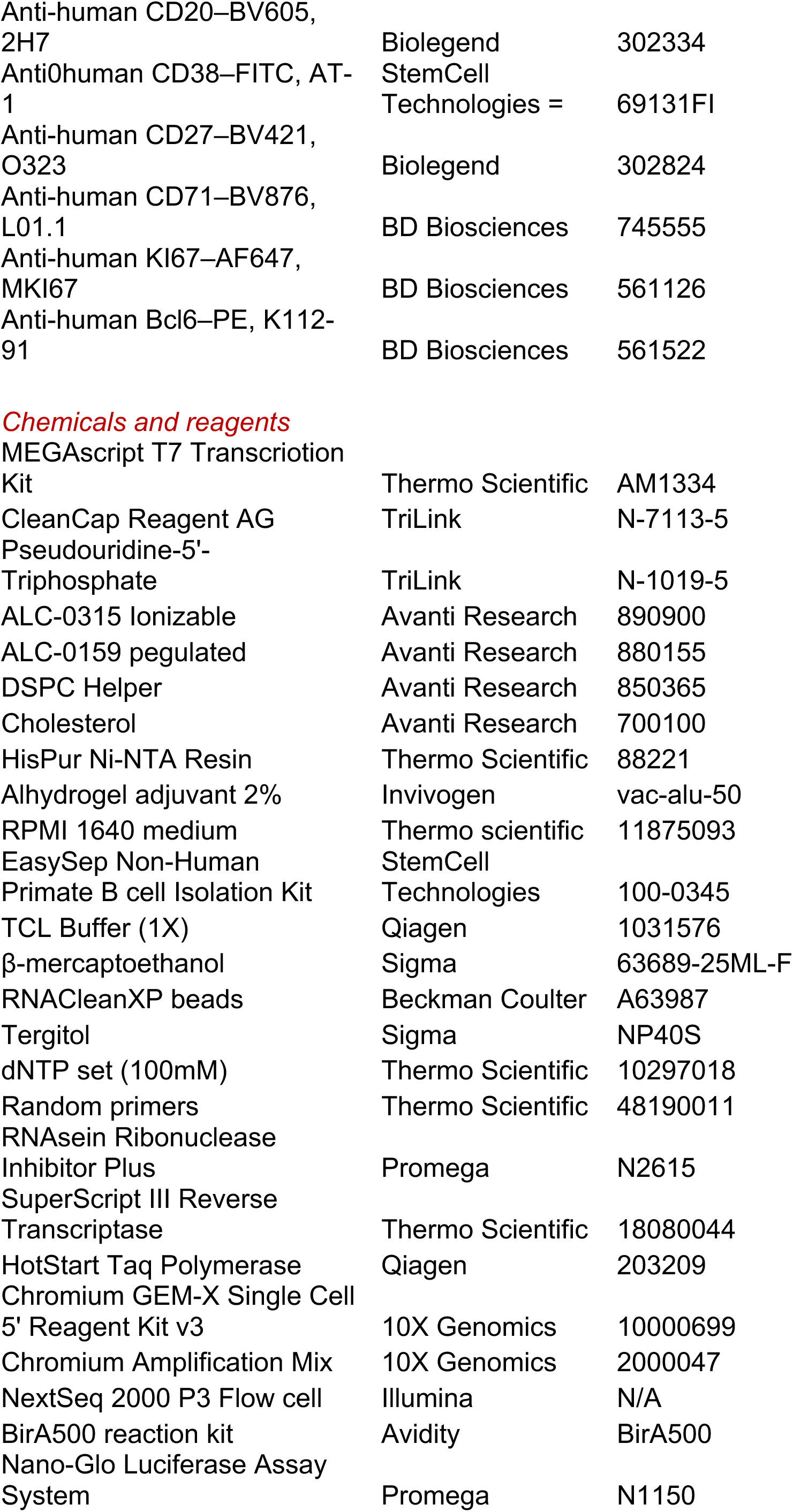
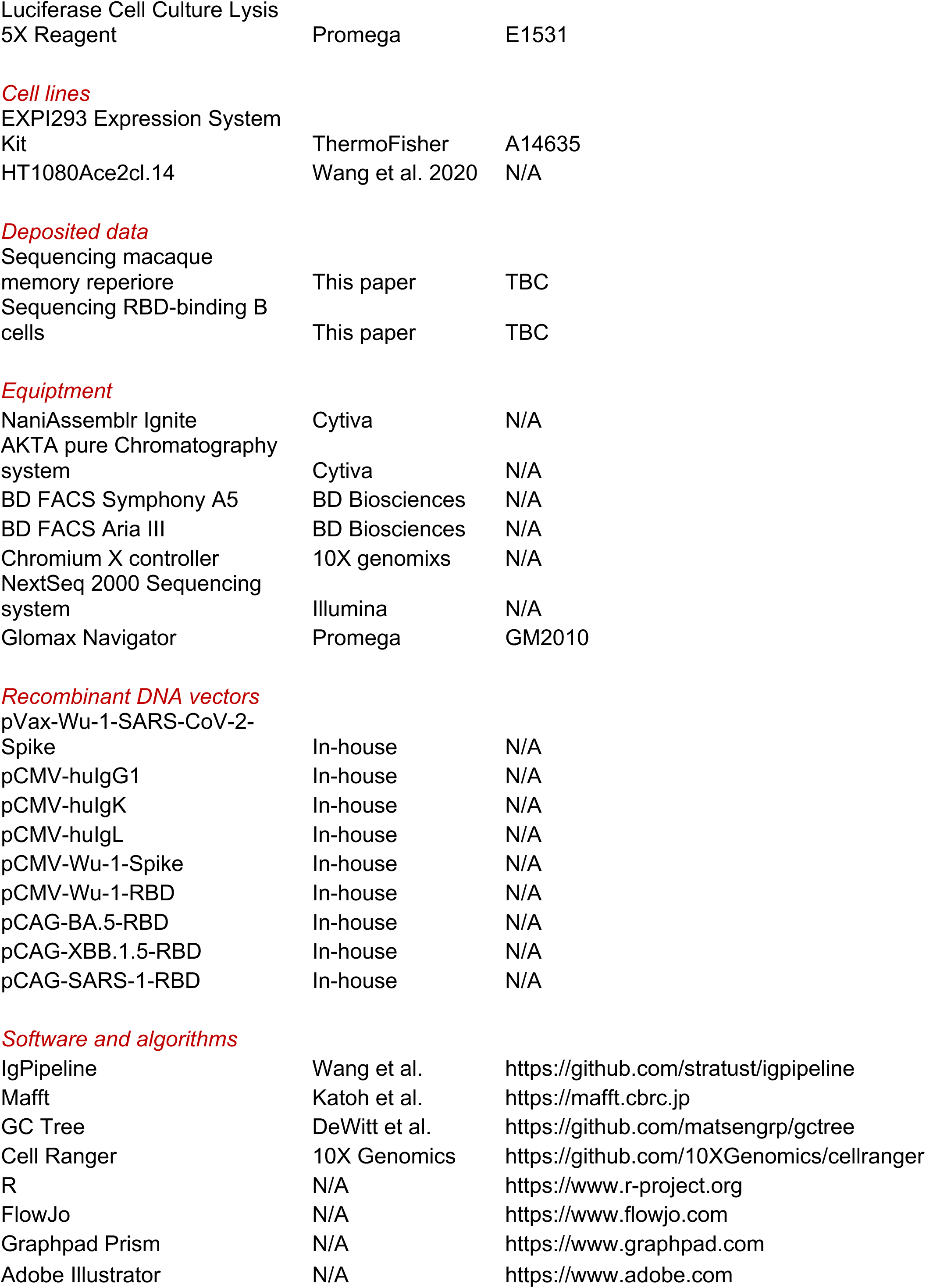

## RESOURCE AVAILIBILITY

### Lead contact

Additional information and requests for resources and reagents should be directed to and will be fulfilled by the lead contact, Michel C. Nussenzweig.

### Materials availability

Reagents, plasmids and mouse lines reported in this study are available upon signing a Materials Transfer Agreement.

### Data and code availability

The latest IgPipeline3 code is publicly available on GitHub (https://github.com/stratust/igpipeline/tree/igpipeline3). Bulk and single-cell RNA-seq are included in both the supplementary tables and are deposited at GEO for access as of the date of publication. Accession numbers are listed in the key resources table. Additional information regarding analysis may be directed to the lead contact.

## EXPERIMENTAL MODELS AND SUBJECT DETAILS

### Rhesus macaques

Six male rhesus macaques (*Macaca mulatta*; Table S1) of Indian genetic origin were maintained in accordance with the Guide for the Care and Use of Laboratory Animals Report number NIH 82-53 (Department of Health and Human Services, Bethesda, Maryland, 1985) and housed in a biosafety level 2 facility. All animal procedures and experiments were performed according to animal study protocols approved by the Institutional Animal Care and Use Committee of National Institute of Allergy and Infectious Disease, NIH. The experimental schedules are outlined in the main text. Briefly, animals were immunized in the deltoid muscle with the standard dose (30 µg mRNA) of the available Pfizer-BioNTech SARS-CoV-2 mRNA vaccine (2023–24 formulation). Auxiliary lymph node clusters, inguinal nodes and the spleen were surgically removed during the necropsies.

### Mice

Six-week-old sex-matched mice were used for all immunizations. Animals were housed in a specific pathogen-free facility with *ad libidum* chow and water. All procedures were performed in accordance with the protocols approved by the Rockefeller University Institutional Animal Care and Use Committee (IACUC). Specific immunization protocols are outlined in the main text. Briefly, mice were immunized with LNPs, where each dose comprised 0.5 µg mRNA contained in LNPs formulated in a low volume (∼15 uL) in the gastrocnemius muscle under isoflurane anaesthesia. Mice were immunized with 2 µg OVA (Sigma) precipitated in 1% Alhydrogel (Invivogen) and administered in the footpad also under anaesthesia.

## METHOD DETAILS

### mRNA–LNPs

For experiments in mice, codon-optimised hexaproline-stabilized (S6P) Wu-1 SARS-CoV-2 Spike ^41^ was cloned into a mammalian *in vitro* transcription mRNA vector. mRNA was synthesised and purified using the MEGAscript T7 Transcription Kit according to the manufacturer’s instructions, except that uridine was substituted for pseudouridine-5’-triphosphate, ^42^ and co-transcriptional capping reagent, CleanCap, was added during the synthesis reaction. mRNA was purified using lithium chloride, washed with 70% ethanol, analysed by agarose gel electrophoresis, and stored at - 80°C. mRNA was encapsulated in LNPs using a nanoparticle assembler to rapidly combine a lipid-containing ethanolic mixture with mRNA in an aqueous citrate buffer. The lipid composition contained ionizible ALC-0315, pegulated ALC-0159, DSPC helper and cholesterol (molar ratio: 0.500:0.015:0.100:0.385). All lipids were stored in EtOH under N_2_ gas at -80°C. The assembled LNPs were dialysed into PBS for immunization.

### Recombinant SARS-CoV-2 Spike and RBD

Spike trimers were expressed as C-terminally His-tagged soluble native-like pre-fusion stabilized trimers.^41^ RBD subunits (GenBank, MN985325.1; Spike protein residues 319–539) were expressed with a C-terminal His-AVI dual tag. Wu-1, BA.5, XBB.1.5 and SARS-CoV-1 RBD variants were cloned into either pCMV or pCAG mammalian expression vectors. Proteins were expressed by transient transfection using the EXPI293 expression system (Life Technologies). Proteins were purified using immobilized metal affinity followed by size-exclusion chromatography (SEC).^20^

### ELISA

ELISAs were performed as described previously.^20^ Serum samples were assayed at a starting concentration of 1:100 and serially-diluted 3–5-fold. Rhesus macaque IgG was detected using the cross-reactive anti-human IgG-HRP secondary antibody used elsewhere.^43^ Monoclonal antibodies were evaluated from a starting concentration of 5 µg/mL.

### Tissue processing

Blood, lymph node and spleen samples were obtained from immunized macaques. PBMCs were isolated from blood using density centrifugation with Histopaque-1077. Single cell suspensions were generated from lymph nodes: nodes were excised, fat was trimmed to expose lymph facia, and were passed through a 100 µm cell strainer before repeated washing with cRPMI. For splenic single cell suspensions, the spleen was first mechanically dispersed before being passed through a 100 µm cell strainer. Red blood cells were lysed using ACK lysis buffer (Gibco) and cells were subsequently washed repeatedly using cRPMI. PBMCs and cell suspensions were transferred into fetal bovine serum (FBS) supplemented 10% DMSO and immediately frozen in a controlled capsule by -1°C/min until samples reached -80°C where they were maintained for at least 24 h prior to LN_2_ transfer until use.

### Flow cytometry and single B cell sorting

Upon thawing cell suspensions, cells were washed with RPMI 1640 medium and B cells were enriched using EasySep Non-Human Primate B Cell Isolation Kit (StemCell Technologies). Cells were incubated in FACS buffer—1X phosphate-buffered (PBS) supplemented with 2% fetal bovine serum (FBS)—with mouse or human Fc block and LIVE/DEAD Fixable near-IR (L/D nIR) for 30 mins on ice prior.

Wu-1 SARS-CoV-2 Spike or RBD tetramers were prepared. First purified recombinant AVI-tagged protein was biotinylated using the BirA500 biotinylation kit according to the manufacturer’s instructions. The biotinylated product was mixed with fluorophore- conjugated streptavidin at a 4:1 molar ratio in PBS at room temperature for 30 mins. Dual two-colour bait staining was used to discriminate specific cells by combining tetramers with the mouse or macaque antibody cocktail indicated below to a final concentration of 10 µg/mL per tetramer. Macaque cells were stained with anti-human CD14–AF700 (clone: 6103), anti-human CD16–AF700 (CB16), anti-human CD3– AF700 (OKT3), anti-human CD8–AF700 (OKT8), anti-human CD20–BV605 (2H7), anti-human CD38–FITC (AT-1), anti-human CD27–BV421 (O323), anti-human CD71– BV786 (L01.1), and PE- and AF647-conjugated RBD tetramers. Mouse cells were stained with anti-mouse CD4–AF700 (GK1.5), anti-mouse CD8–AF700 (53-6.7), anti- mouse F4/80–AF700 (BM8), anti-mouse Gr-1–AF700 (RB6-8C5), anti-mouse B220– eFluor450 (RA3-6B2), anti-mouse CD38–PE/Cy7 (90), anti-mouse Fas–PE (SA367H8), anti-mouse CD138–FITC (281-2), and BV605- and AF647-conjugated RBD tetramers. The macaque staining cocktail was prepared in FACS buffer supplemented with Brilliant Stain Buffer Plus.

Cell sorting was performed using a FACS Aria III. Memory (L/D nIR^-^CD14^-^CD16^-^CD3^-^ CD8^-^CD20^+^CD38^+^CD27^+^) and GC (L/D nIR^-^CD14^-^CD16^-^CD3^-^CD8^-^CD20^+^CD38^lo^CD71^hi^) B cells were isolated from macaque cell suspensions. Gating strategies are shown in the results. Cells used for single cell mRNA sequencing using the 10X Chromium platform were sorted in bulk into PBS with 0.5% FBS for immediate use. Single cells were sorted directly into 96-well plates containing a reverse transcriptase reaction cocktail described below.

### Single B cell sequencing and cloning

For sorted single macaque cells, cDNA was synthesised by reverse transcription with SuperScript III Reverse Transcriptase in a reaction mixture containing 8 ng/µL random primers and 0.6 U/µL of RNase inhibitor. cDNA was stored at -20°C until subsequent polymerase chain reaction (PCR) steps.

Antibody gene amplicons were synthesised by nested PCR using the primers in table S5.^44,45^ All PCRs comprised of 50 elongation cycles of 55 s. Macaque annealing temperatures (*IgH/IgK/IgL*): first PCR, 55/51/51; second PCR, 60/58/58. Amplicons were confirmed by gel electrophoresis and Sanger sequenced (Genewiz) using the respective reverse primer(s) from the second-round PCR.

Heavy/light chain variable region gene fragments were synthesised (IDT) and cloned into linearised human Ig expression vectors (NCBI GenBank accession numbers FJ475055, FJ475056 and FJ517647) using Gibson assembly ^20^.

### RNA sequencing (10X)

Sorted macaque B cells were loaded onto the Chromium X controller for bead formation using the Chromium Next GEM Single Cell v3.1 kit. Cell-barcoded cDNA libraries were generated according to the manufacturer’s instruction. VDJ libraries were prepared using the Chromium Single Cell VDJ enrichment mix, using custom primers.^13^ Libraries with unique Illumina adapters were pooled and sequenced using a NextSeq 2000 sequencing system.

### Antibody production and purification

Recombinant antibodies were expressed by transient transfection in EXPI293 cells and purified from supernatant using Protein G agarose resin.^20^ Bulk anti-S6P Ig was purified from pooled serum in immunized mice and the mock Ig from naïve mice. Ig was purified Protein G.

### In vitro neutralization assay

SARS-CoV-2 pseudotyped particles were generated as described.^20,46^ Briefly, 293 T cells were transfected with pNL4-3ΔEnv-NanoLuc and pSARS-CoV-2 Spike_Δ19_. Particles expressing Wu-1 or select variants were produced and stored at -80°C.

Serially diluted plasma samples or monoclonal antibodies were pre-incubated for 1 h with pseudovirus prior to incubating with HT1080Ace2cl.14 cells for 48 h. Cells were lysed with Luciferase Cell Culture Lysis reagent. Luciferase activity was measured with the Nano-Glo Luciferase Assay system, and data were acquired using the CLARIOstar Plus reader. Data were normalised to the negative control wherein no plasma/monoclonal antibody was added. Each test was conducted in duplicates.

Neutralisation titres were determined using a four-parameter non-linear regression (constraints: top = 1, bottom = 0).

## QUANTIFICATION AND STATISTICAL ANALYSIS

### Single-cell library processing

To increase the limited repertoire of V genes for rhesus macaques available in the IMGT database, we compiled a custom germline sequence database that amalgamates the IMGT database (as updated Jan 2025) ^47,48^ and the private rhesus macaque database previously published and kindly shared by Dr Shane Crotty’s group.^12,13^

Hashtag-oligos unique molecular identifier quantification were performed with Cell Ranger multi v.8.0.1 (10X Genomics), using the Cell Ranger GEX reference Mmul_8.0.1, and analyzed in R with Seurat v.5.1.0.^49^ Cells were demultiplexed with MULTISeqDemux, and those classified as doublets were excluded. Single-cell BCR libraries processed with Cell Ranger VDJ v.8.0.1 without a VDJ reference genome using the parameters *–denovo –chain=auto* and subsequently annotated with igblastn v.1.22.0 using our custom Rhesus macaque germline database described above. Contigs containing fewer than 50 reads and more than one heavy or light chain were removed.

### Computational analyses of antibody sequences

Heavy and light chains of both single-cell BCR and Sanger reads were paired and analyzed using IgPipeline v.3.0 ^50^ using the custom rhesus macaque germline gene segment database described above. Scripts for sequence annotation, processing and graphic rendering are publicly available on GitHub (https://github.com/stratust/igpipeline/tree/igpipeline3).

The paired IgH and IgL chains of antibody sequences inferred as identical clonal progeny were merged and aligned to the cognate germline sequence in the custom database described above, using mafft v.7.520 ^51^ with default parameters except for *--globalpair*. Genotype-collapsed phylogenetic trees of clonal lineages were inferred using GCTree v.4.1.2 (https://github.com/matsengrp/gctree).^52^ Each node represents a unique IgH and IgL combination with number within each node indicating the number of identical sequences. The scales represent the branch lengths, estimated based on the number of nucleotide mutations.

### Data processing and statistics

All statistical tests were calculated in Prism 10 or R (v.4.4.1). Specific statistical information, including *n* and statistical significance values, are indicated in the text and figure legends. For log-transformed data, the geometric mean is used to indicate central tendency, unless otherwise indicated; correspondingly, for non-rank-based statistical tests, groups were compared using the log-transformed data. Chao1 diversity estimates were conducted using the iNEXT script.^53^ All comparisons are two-tailed and multiple comparisons are adjusted for false discovery.

## Supplementary Figure Legends

**Figure S1:**
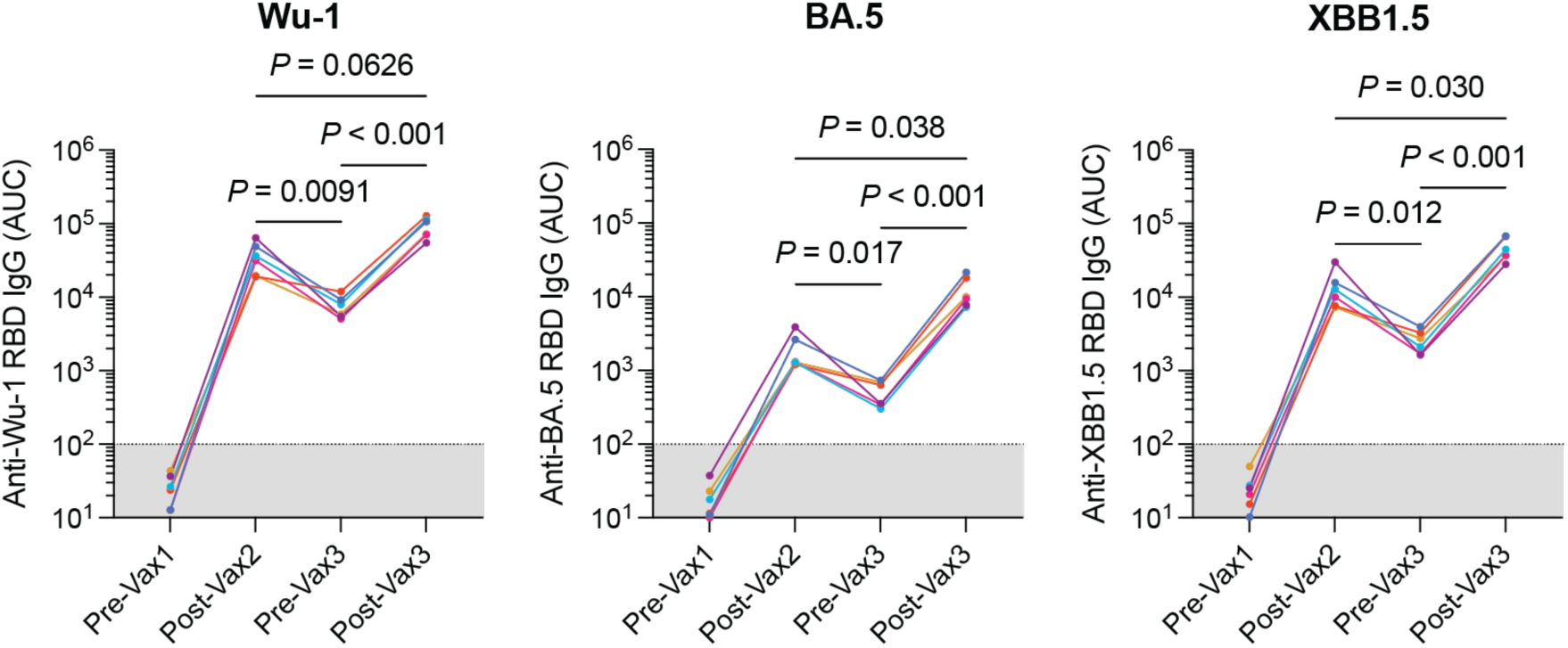
Plasma reactivity against SARS-CoV-2 RBD variants following immunization. Plots showing IgG reactivity of plasma from rhesus macaques iteratively immunized with Pfizer Comirnaty 2023–24. Plasma was tested against RBD variants. Connected dots indicate data from the same animal at different timepoints. Comparisons were conducted using paired Tukey’s post-hoc comparisons.

**Figure S2:**
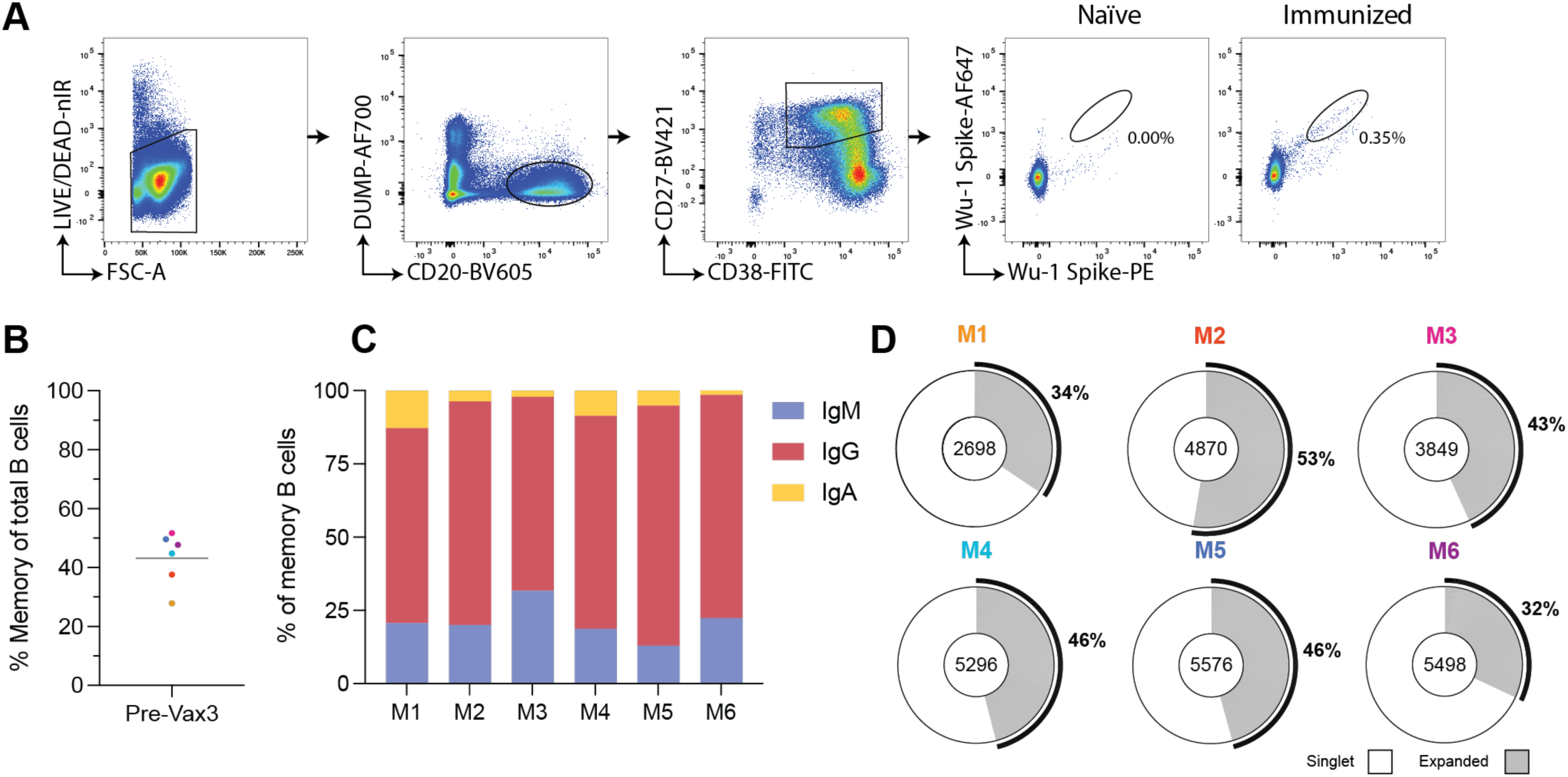
Circulating memory B cell clonality in rhesus macaques. **(A)** Flow cytometry plots showing the gating used to select memory B cells (CD38^+^CD27^+^) in PBMCs and representative bait-binding profile in macaques. **(B)** Plot showing the percentage of circulating memory B cells pre-vax3 in each animal. Each dot is the value from a single animal, colorized according to the macaque ID. Bar denotes the mean. **(C** and **D)** Bulk memory B cell antibody sequences were acquired using the 10X genomics platform. **(C)** Graph shows the distribution of isotypes represented in the memory compartment for each macaque. **(D)** Donut plots showing the distribution of expanded versus singlet memory clones. The number of heavy and light chain variable region pairs obtained from each animal is shown in the middle of the donut. Grey segment and marked percentages reflect the proportion of cells belonging to expanded clones, while white denotes singlets.

**Figure S3:**
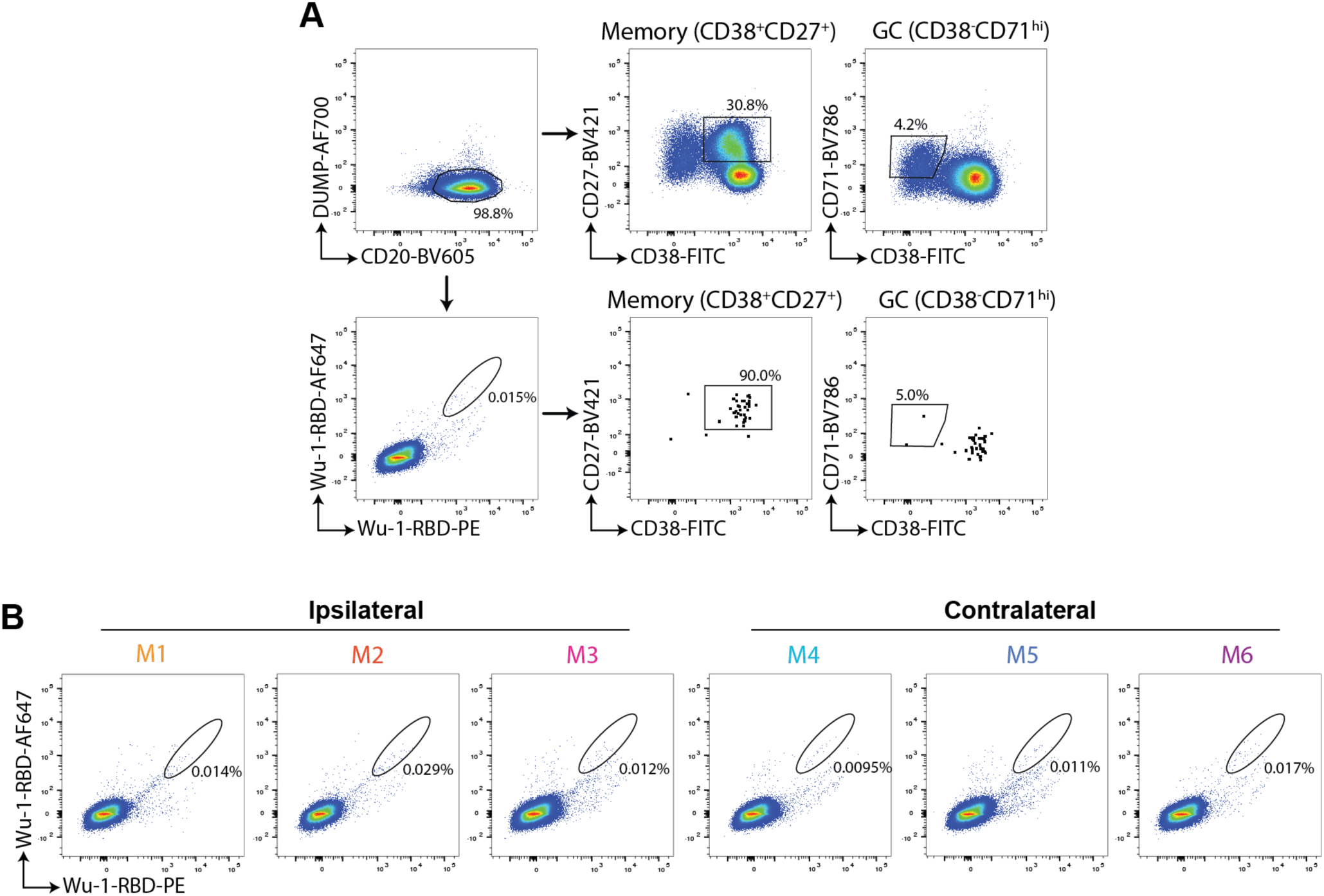
RBD-specific B cells in macaque spleen following booster immunizations. **(A)** Flow cytometry plots of splenic macaque B cells. Plots show back-gating on Wu-1 bait-binding B cells. **(B)** Plots showing the population of splenic memory B cells that bind RBD in each animal. Spleens were harvested at wk 15 following the third immunization.

**Figure S4:**
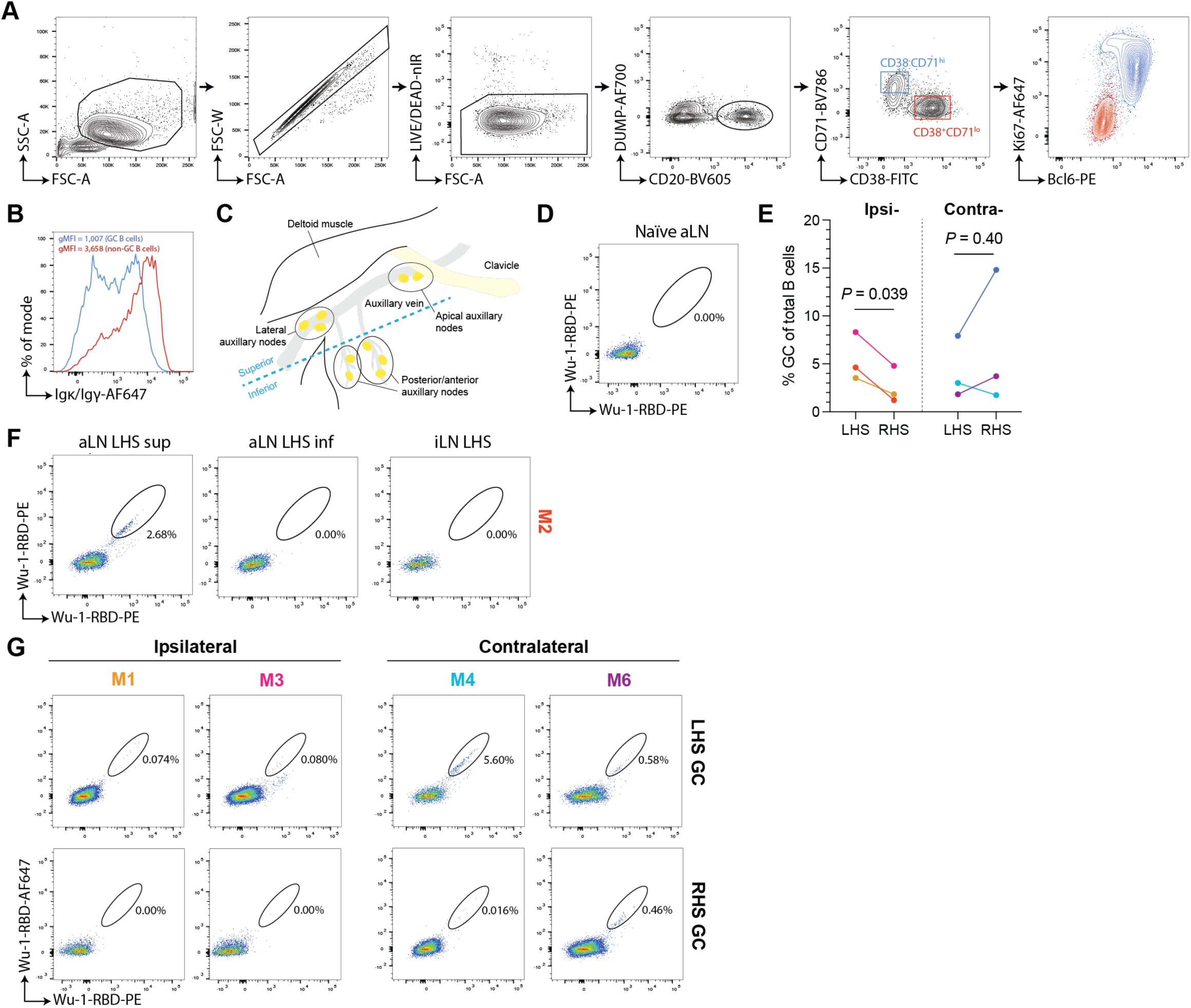
Flow cytometric analysis of booster responses in Rhesus macaques. **(A)** Contour plots showing the lineage markers CD20^+^CD38^-^CD71^hi^ that efficiently identifies GC B cells in the lymph nodes of Rhesus macaques. **(B)** Histogram showing the distribution of surface BCR densities on GC and non-GC B cells. **(C)** Schema highlighting the anatomy of major auxiliary node clusters in a Rhesus macaque. **(D)** Flow cytometry plot of Wu-1 RBD bait-binding on aLN GC B cells from a naïve macaque. **(E)** Plot showing the percentage of GC B cells in the respective superior aLN. Connected dots indicate data from the same animal. Data were compared using paired Student’s t-tests. **(F)** Representative pseudocolour plots showing RBD bait-binding on pre-gated GC B cells isolated from either the superior auxiliary lymph node (sup aLN), the inferior (inf) aLN or the inguinal (i)LNs. **(G)** Flow cytometry plots showing RBD bait-binding on GC B cells in the post-boost sup aLN.

**Figure S5:**
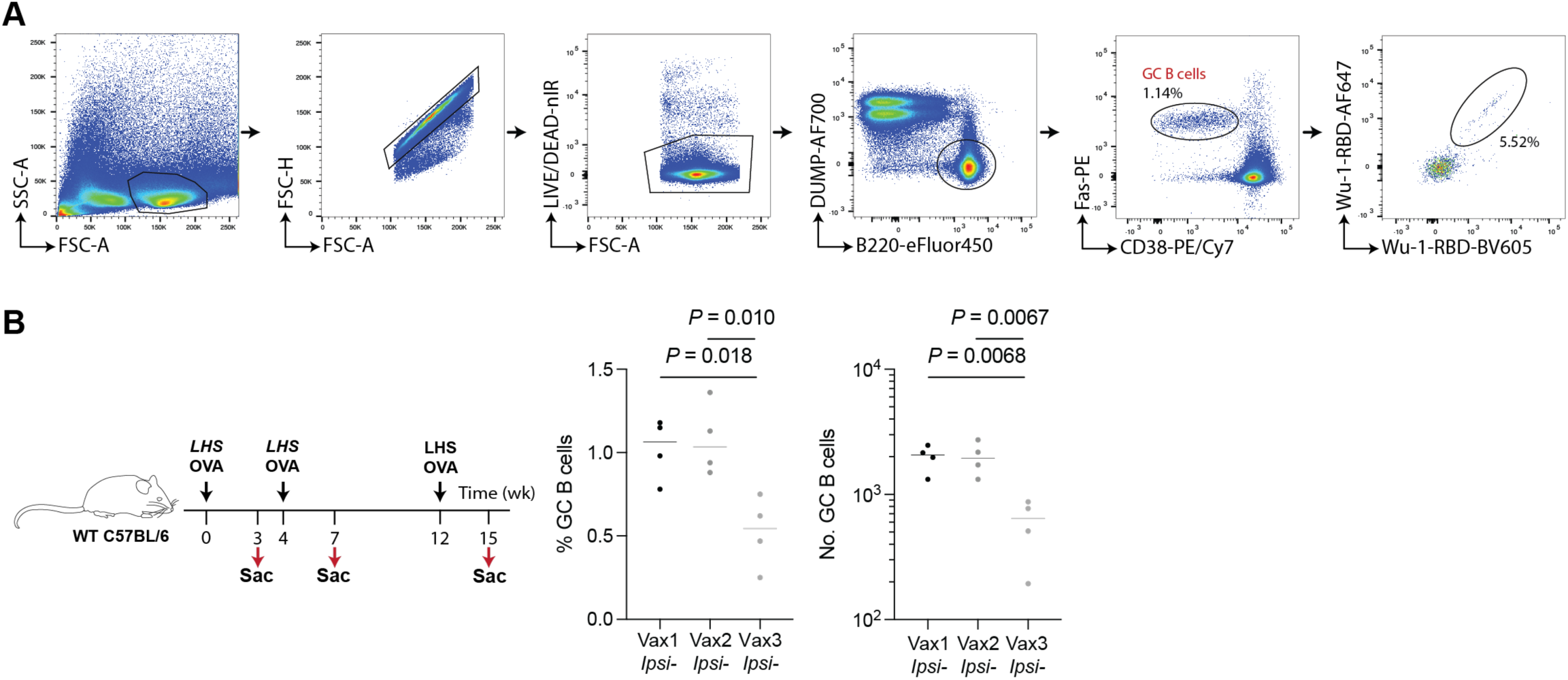
Germinal center B cell reactions following iterative immunizations in mice. **(A)** Mouse GC and RBD-bait binding B cell gating strategy. **(B)** Experimental scheme of immunization schedule (left) and evaluation of GC size after immunization (right).

**Figure S6:**
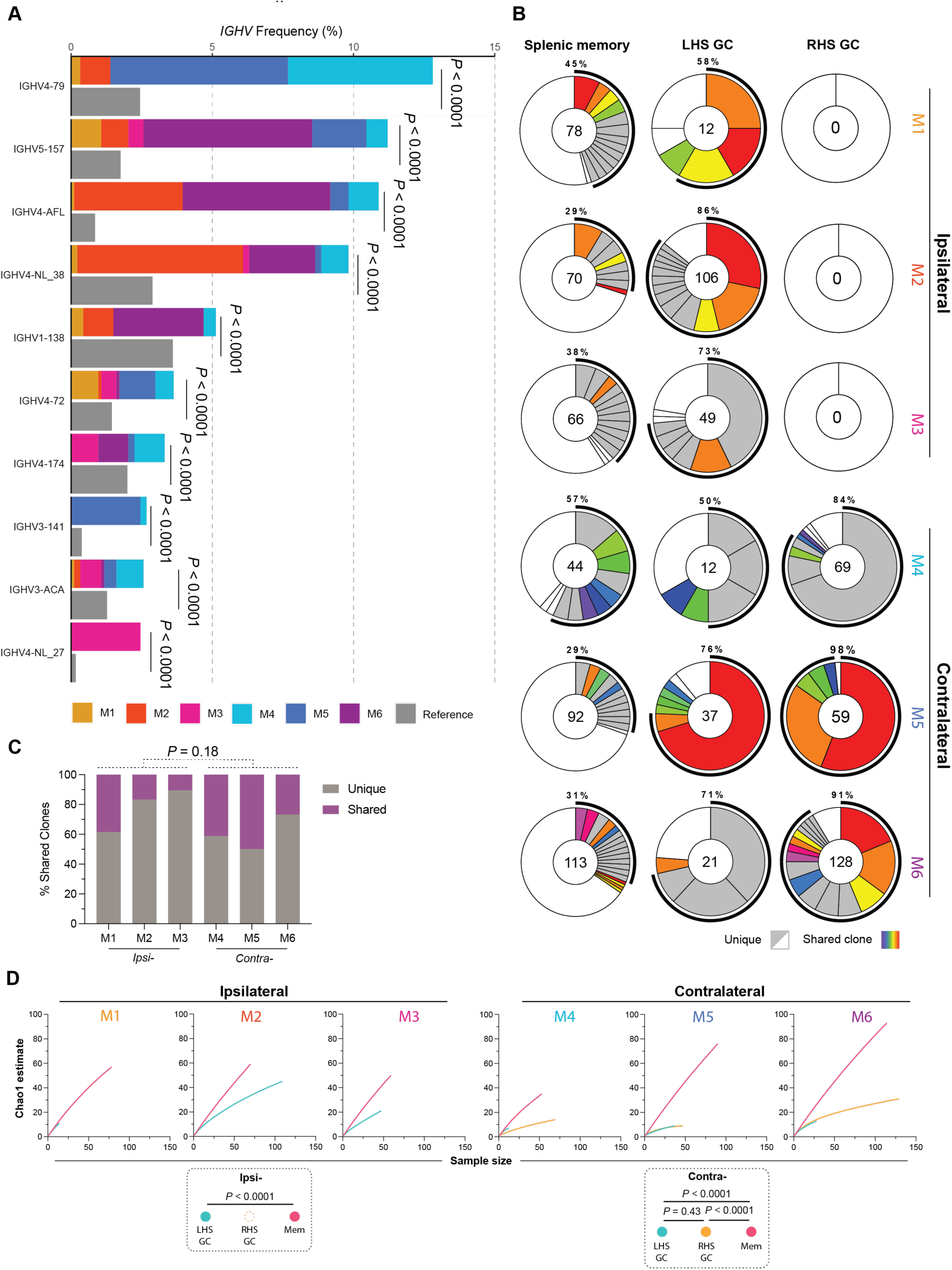
Clone sharing and V_H_ gene distribution of post-boost RBD-binding B cells. **(A)** Bar plot shows the relative abundance of the 10 most common *IGHV* gene segments utilized by all RBD bait-binding B cells sequenced. Bars were partitioned and colorized according to animal identity. This was compared to the bulk circulating memory B cell library (fig. S2; and table S2). Statistical significance was measured using a two-sided binomial test with Benjamini-Hochberg *P*-value correction. **(B)** Donut plots showing the clonality of Wu-1 RBD bait-binding B cells from both memory and GCs. The number of sequences reflected in each plot is shown in the center of each plot. Colored/grey segments denote expanded clones, with their size proportional to the number of clone members. The outer ring segment and marked percentage represents the proportion of cells belonging to an expanded clone. White segments denote singlets. Colored segments for each given animal correspond to clones present across compartments. ND = not detected/no sequences recovered. **(C)** Plot showing the proportion of clonal families shared across multiple compartments or unique to a single compartment. Boost condition effects were tested using a Student’s t test. **(D)** Plots showing the Chao1 estimate traces for both memory and GC B cells. Compartment diversity was compared using a Kolmogorov-Smirnov test.

**Figure S7:**
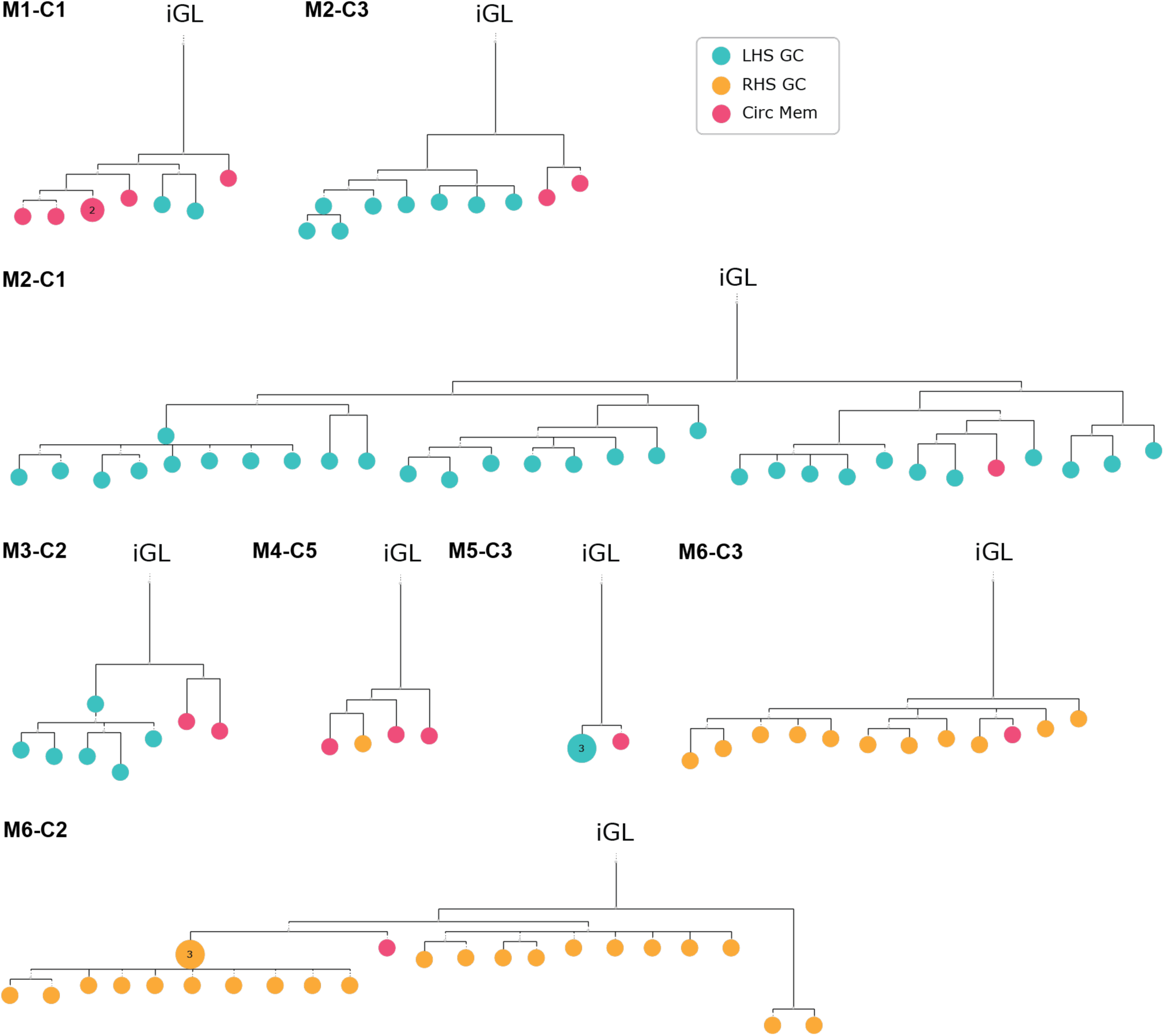
Representative trees of clones shared between the memory and GC compartments. Phylogenetic trees of representative B cell clonal families shared between the GC and memory. Distance is proportional to the heavy and light chain sequence disparity, rooted to their inferred germline revertant (iGL). Unless otherwise marked, all nodes comprise of a single cell. Node color denotes the location and fate of the originating cell.

**Figure S8:**
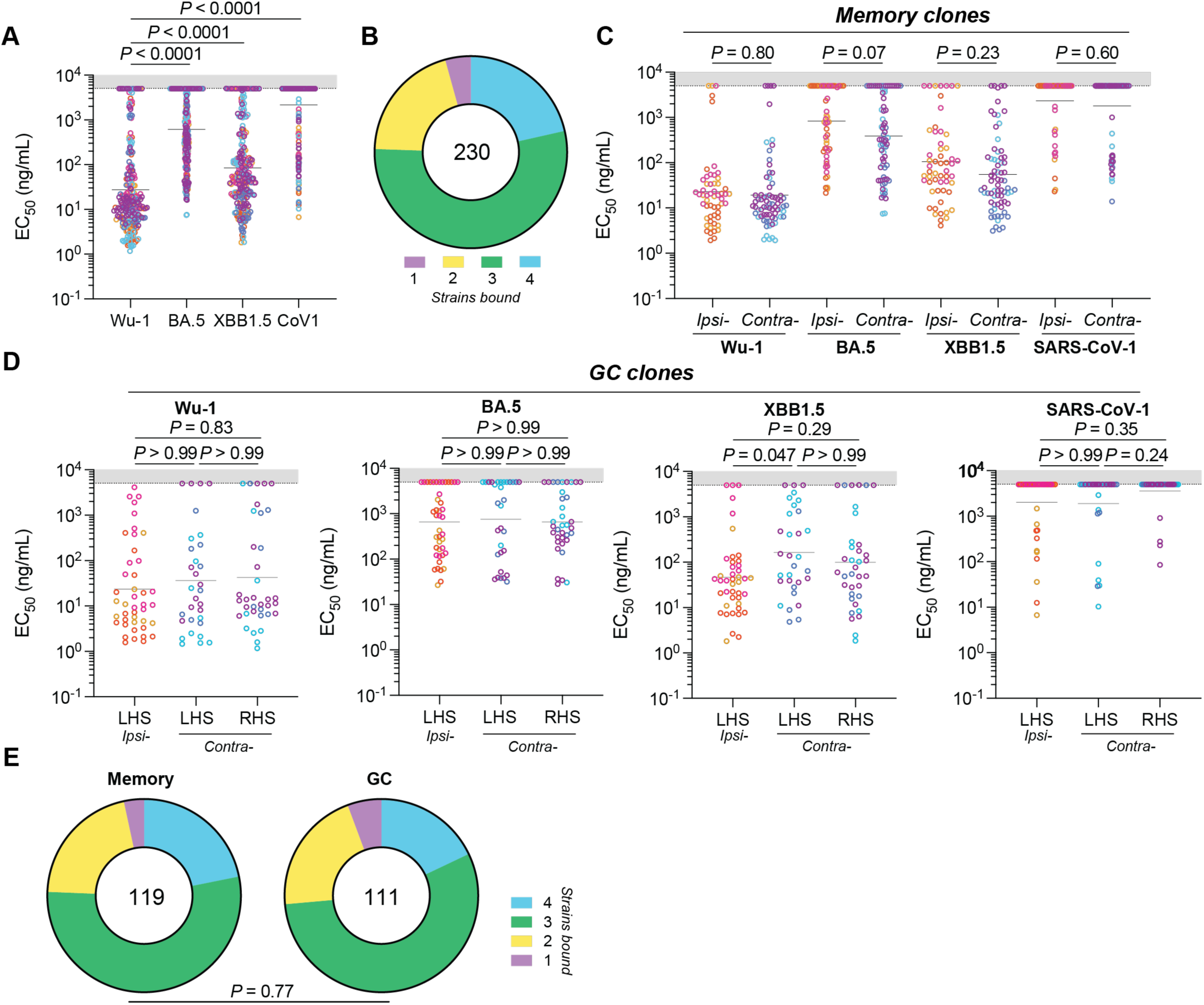
Binding and breadth of anti-RBD monoclonal antibodies. **(A)** Plots showing the EC_50_ values of monoclonal antibodies cloned from memory and GC B cells screened against RBD variants via ELISA. The total number of monoclonal antibodies tested was 247. **(B)** Donut plot showing the cross-variant RBD binding for antibodies reactive with Wu- 1 RBD. Strains were deemed binders if EC_50_ < 5 µg/mL. **(C)** Graph showing RBD variant binding of antibodies cloned from memory B cells in both ipsilaterally and contralaterally boosted animals. Vax3 site conditions were compared using Mann-Whitney tests. **(D)** Plots show RBD binding of antibodies cloned from GC B cells. **(A** and **D)** Groups were compared using Kruskal-Wallis tests. **(A, C and D)** Dots show the EC_50_ value of a monoclonal antibody an indicated RBD variant. Dots are colorized according to the animal an antibody was cloned from. Bars denote the geometric mean. **(E)** Donut plots showing the RBD reactivity breadth of antibodies derived from memory or GC B cells. Groups were compared using a chi-squared test. **(B and E)** Clones where Wu-1 RBD EC_50_ ≥ 5 000 ng/mL was excluded from this analysis.

**Figure S9:**
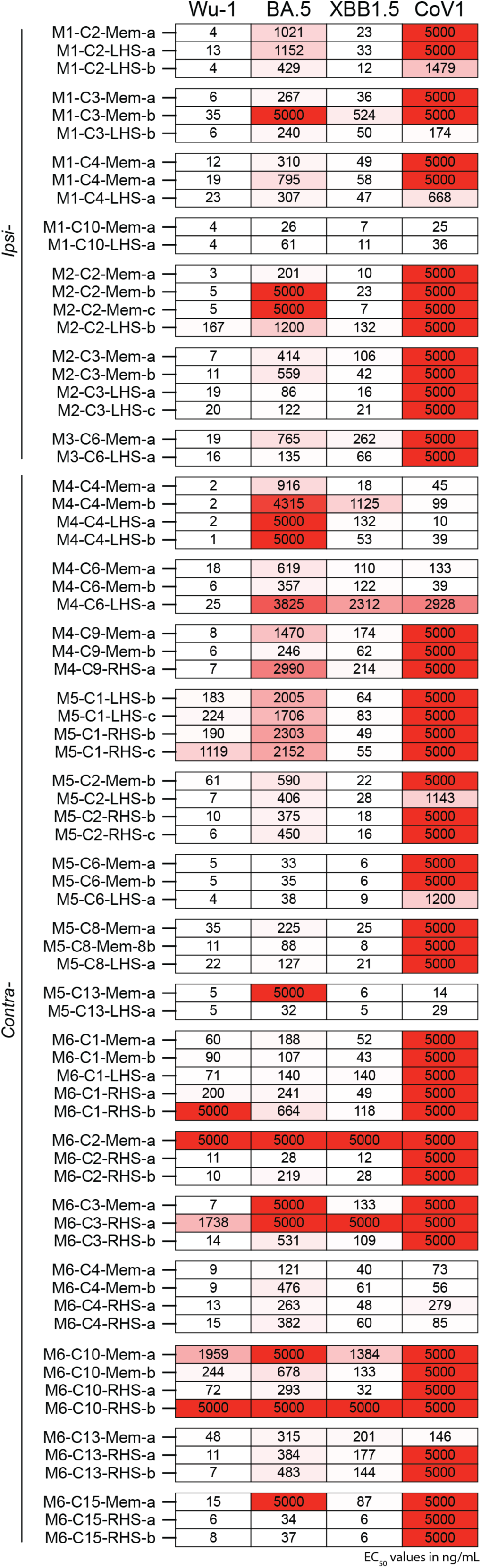
Variation in RBD reactivity of monoclonal antibodies via ELISA. Heatmap showing the EC_50_ values of monoclonal antibody reactivities against RBD variants via ELISA. Cells are clustered by shared clones, and thus antibodies derived from cells present in multiple compartments were evaluated. mAb ID assignments correspond to the monkey (M*x*), clone number (C*y*), origin (memory, LHS/RHS GC) and cell identifier (a,b, etc.).

## Supplementary Table Legends

**Table S1**

Individual Rhesus macaque characteristics.

**Table S2**

Sequence library of bait-agnostic macaque circulating memory B cells.

**Table S3**

Sequences of RBD bait-binding heavy and light chain pairs from macaque memory and GC B cells.

**Table S4**

Sequences, EC_50_ values and NT_50_ values of cloned monoclonal antibodies.

**Table S5**

Macaque primers used for single-cell BCR amplification and sequencing.

